# Amphisome biogenesis couples presynaptic autophagy to local protein synthesis

**DOI:** 10.1101/2025.09.23.678051

**Authors:** Maria Andres-Alonso, Catrin Schweizer, Carolina Montenegro-Venegas, Sarah Wirth, Carola Schneider, Rabia Turacak, Katarzyna M. Grochowska, Rabia Bice Aydin, Lisa Mahnke, Shuting Yin, Anna Karpova, Rudolph Reimer, Antonio Virgilio Failla, Tobias M. Boeckers, Eckart D. Gundelfinger, Michael R. Kreutz

## Abstract

Synaptic neurotransmission imposes high demands on membrane turnover, metabolism, and the remodeling of presynaptic molecular composition. While the impact of autophagy on neurotransmission has been firmly established, evidence for activity-dependent synaptic induction of autophagy remains surprisingly limited. Here, we demonstrate that amphisomes containing BDNF/TrkB are formed at presynaptic boutons following sustained synaptic activation. Activity-dependent bulk endocytosis serves as a membrane source for amphisome biogenesis, while key autophagy proteins are recruited to the active zone, and autophagy initiation is triggered locally by the energy-sensing kinase AMPK. BDNF/TrkB-containing amphisomes contribute to the turnover of key presynaptic cytoskeletal proteins involved in synaptic vesicle clustering. The formation of amphisomes following sustained synaptic activity facilitates both the degradation of these proteins and their replenishment through local translation of their mRNAs at presynaptic boutons. We propose that activity-induced synaptic autophagy largely reflects amphisome formation, which in turn is required for the replacement of proteins within the local presynaptic cytomatrix.

## Introduction

Neurons are postmitotic cells characterized by long processes and a large number of synaptic contacts located far away from the soma. The extreme length and narrow caliber of axons pose significant challenges for proteostasis at presynaptic boutons; however, the mechanisms that ensure efficient protein replenishment in these compartments remain poorly understood. In particular, little is known on how scaffold proteins are targeted to degradation at boutons without compromising presynaptic function. The presence of mRNA and ribosomes as well as compelling evidence for local protein translation points to the possible involvement of activity-dependent mechanisms in local exchange of proteins at boutons^1–3^. Notably, the presynapse contains densely packed protein scaffolds that are essential for synaptic neurotransmission^4–8^. The organization of presynaptic compartments through liquid–liquid phase separation has emerged as a key principle underlying processes such as the spatial localization and short-range trafficking of synaptic vesicles (SVs) within vesicle clusters^9, 10^. These biomolecular condensates are composed of highly abundant protein components including Synapsin, Tau and those of the cytomatrix of the active zone (CAZ).

Macroautophagy (hereafter referred to as autophagy) is a major degradative pathway, tailored to the disposal of large protein complexes, toxic aggregates, and even entire organelles^11^. In contrast to non-neuronal cells, autophagy in neurons is highly compartmentalized. Under basal conditions, autophagosome formation occurs predominantly in axons, from where autophagosomes are transported retrogradely to the soma, which contains the majority of degradative lysosomes^11–14^. Phagophore formation in axons under basal conditions crucially involves the endoplasmic reticulum (ER) and ER-derived membranes^15^. Although other membrane sources have been proposed it is likely that neuron-specific adaptations in the mechanisms of phagophore formation exist and that synaptic activity is a key determinant in this process^11, 16^. However, our understanding about the induction of autophagy following synaptic activity, its function, and the underlying mechanisms of activity-dependent synaptic autophagy is currently sparse.

Intricate mechanisms are in place to ensure synaptic neurotransmission with high fidelity and the consequences of autophagy for the functional integrity of presynaptic boutons can be potentially detrimental. How is autophagy triggered and controlled in compact presynaptic compartments of mammalian neurons? We here show that in hippocampal neurons sustained neurotransmission leads to autophagy activation at presynapses which crucially results in the formation of TrkB-amphisomes. Biogenesis of amphisomes depends upon bulk endocytosis and the recruitment as well as activation of key autophagy proteins, like ULK1, induced by the energy-consumption sensing enzyme 5’-AMP-activated protein kinase (AMPK)^17–19^. Signaling of high-energy demand via AMPK during sustained SV release and synaptic membrane retrieval converge to generate signaling amphisomes that engulf key cytomatrix proteins of the presynapse. Signaling BDNF/TrkB-amphisomes associate with ribosomes and TrkB-signaling from amphisomes induce the local translation of cytoskeletal presynaptic components of the SV cluster and the CAZ.

## Results

### Amphisome biogenesis is coupled to synaptic activity

Amphisomes are organelles of the autophagy pathway that result from fusion of late endosomes to autophagosomes. At present it is debated whether amphisomes in neurons serve signaling function by incorporating active TrkB receptors^20, 21^ or on the contrary that TrkB predominantly traffics in signaling endosomes^22, 23^. We reasoned that biogenesis of TrkB-signaling amphisomes might be linked to synaptic function and that neurotransmission in mature neurons might underlie their biogenesis in an activity-dependent manner. To test this hypothesis, we first assessed the presence of TrkB-containing amphisomes in axons from rat hippocampal neurons (DIV 5 and DIV 7) before or during early synaptogenesis in comparison to differentiated neurons (DIV 14-15) that maintain several thousand functional synaptic contacts (Extended Data Fig. 1a-b). We indeed found that phosphorylated TrkB (pTrkB) extensively co-localized with LC3-positive vesicles in axons immunopositive for Tau at DIV 14 (Fig. 1a-e, Extended Data Fig. 1c). However, we did not observe co-localization at DIV5 (Fig. 1a-e, Extended Data Fig. 1c) whereas with the onset of synaptogenesis at DIV7 we detected a slight but still non-significant increase in LC3 vesicles positive for pTrkB (Extended Data Fig. 1c). The density of LC3-positive vesicles was significantly increased in neurons at DIV14 (Fig. 1c) and in differentiated neurons more than one third of autophagic vesicles was positive for active TrkB receptors (Fig. 1d). Correspondingly, more than 40% of immunofluorescent pTrkB puncta colocalized with LC3 (Fig. 1e).

**Figure 1.**
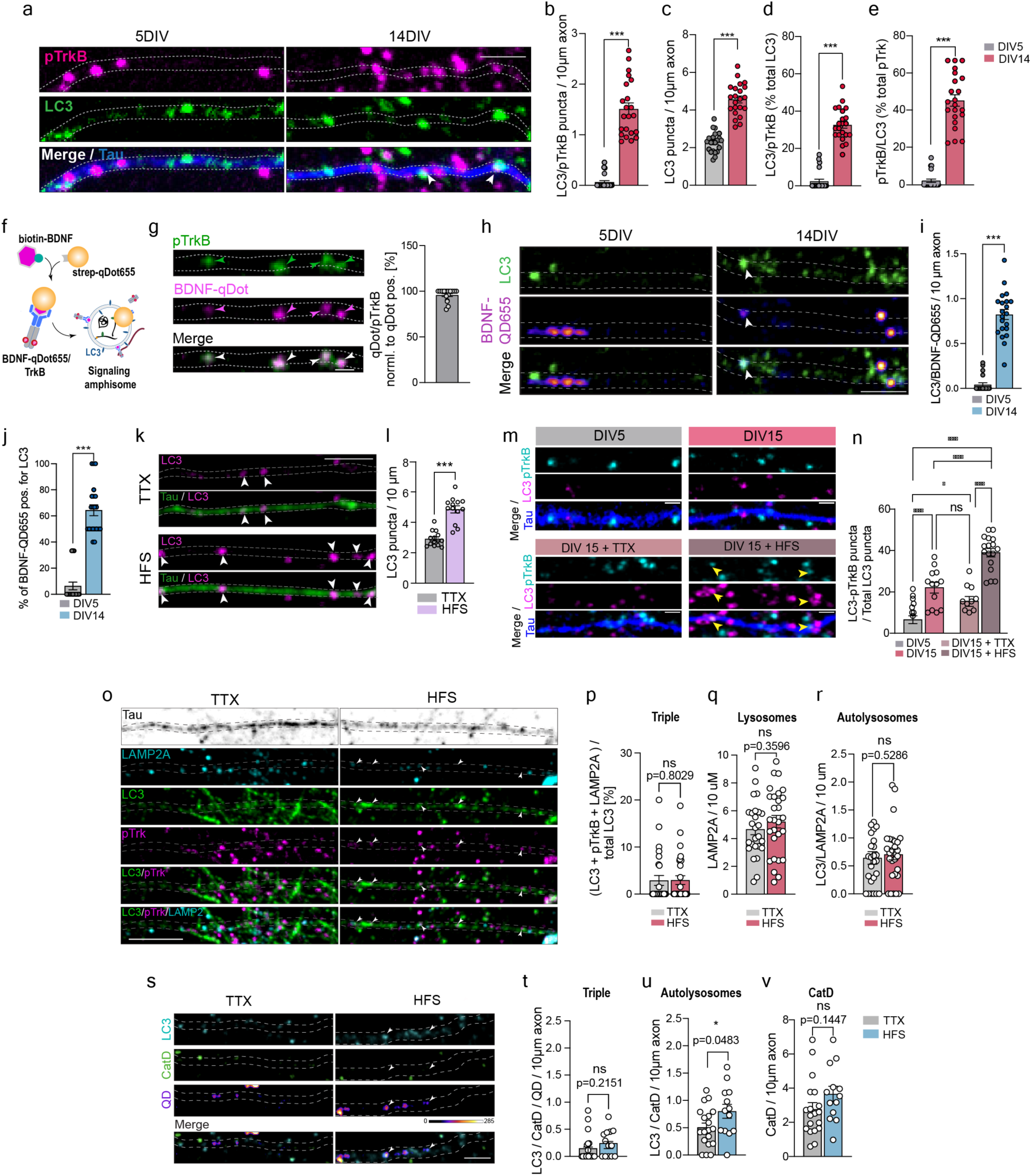
Signaling amphisomes are segregated from the endo-lysosomal pathway in mature neurons. A) Representative images of axons identified by Tau from primary hippocampal neurons at DIV5 and 14 immunostained against pTrk and LC3. Scale bar indicates 2µm. B) Number of LC3- and pTrkB- double positive puncta in 10 µm axon of neurons at DIV5 and DIV14 from the experiment in A. C) Bar graph depicting the number of LC3-positive puncta per 10µm of axon obtained from neurons at DIV5 and DIV14 from the experiment in A. D) Quantification of amphisomes (LC3/pTrkB-positive) in 10µm axon as percentage of the total LC3 from axons at DIV5 and DIV14. E) Percentage of all pTrkB puncta that colocalize with LC3 puncta in 10µm axon axons at DIV5 and DIV14. (B-E) *n=22 from 3 independent cultures* F) Schematic representation of the assay employed to label BDNF/TrkB complexes using biotin-BDNF coupled to streptavidin-qDot655. G) Representative images of primary neurons incubated with BDNF-qDot655 and immunostained against pTrkB. Scale bar indicates 1µm. Right, bar graph showing the percentage of BDNF-qDot655 positive for pTrkB in axons labeled by Tau (not shown, shaded line). *(n=16 from 3 independent cultures)* H) Representative images of endogenous LC3 staining and BDNF-qDot655 in axons (depicted with scattered white lines) from primary hippocampal neurons at DIV5 and 14. Scale bar indicates 2µm. I) Bar graph shows colocalization of LC3 puncta with endogenous BDNF-qDot655 per 10µm axon in DIV5 compared to DIV 14. Extracellular signal of qDot655 was quenched. J) Quantification of axonal BDNF-qDot655 positive for LC3 puncta as percentage. (G-H) *n=21 from 3 independent cultures* K) Representative image of LC3 puncta in axon labeled by Tau in TTX and HFS-stimulated cells. Scale bar indicates 2µm. L) Quantification of experiment in K) showing the amount of LC3 puncta per 10µm axon. *n=13 (FS) n=14 (TTX) from 3 independent cultures* M) Representative images of primary neurons immunostained for LC3, pTrkB and Tau in neurons at DIV5 vs. DIV15 neurons under basal conditons as well as DIV15 neurons that were silenced with TTX or stimulated using high-frequency stimulation (HFS). Scale bar indicates 1µm. N) Quantification of amphisomes (pTrkB/LC3-positive) in axons as percentage of total LC3-positive vesicles from the experiment in M. n=*3 independent cultures)* O) Representative images of endogenous LC3, pTrkB and LAMP2 staining in axons from primary hippocampal neurons at DIV14 comparing TTX and HFS conditions. Scale bar indicates 2µm. P) Quantification of triple-positive structures (pTrkB/LC3/LAMP2) as percentage of total analyzed LC3. Q) Quantification of LAMP2A puncta in 10 µm axon. R) Bar graph shows colocalization of LC3 puncta with LAMP2A puncta per 10 µm axon. S) Representative images of endogenous Cathepsin D staining and BDNF-Qdots 655 in axons from primary hippocampal neurons at DIV14 comparing TTX and HFS conditions. Scale bar indicates 2µm. T) Bar graph shows colocalization of Cathepsin D and LC3 puncta with endogenous BDNF-QD655 per 10µm axon in DIV 14. Extracellular signal of qDot655 was quenched. U) Bar graph shows colocalization of LC3 puncta and Cathepsin D puncta in 10µm axon. V) Quantification of Cathepsin puncta per 10µm axon. (M-P) *n=26 (HFS) n=28 (TTX) from at least 3 independent cultures* (P-S) *n=13 (HFS) n=19 (TTX) from at least 3 independent cultures* Data are plotted as average ± SEM. ^∗∗∗^ represents p < 0.0001 to 0.001 by two-tailed Mann-Whitney U test.

We next visualized BDNF internalization by taking advantage of BDNF coupled to qDots655 (Fig. 1f). BDNF-qDots655 was added to the medium and the use of a non-cell-permeable fluorescence quencher allowed us to identify internalized qDots that were consistently positive for pTrkB indicating that they can serve as marker for signaling amphisomes (Fig. 1g). We indeed confirmed that BDNF/TrkB complexes prominently localize to LC3-positive autophagosomes at DIV14, but not at DIV5 (Fig. 1h-j; Extended Data Fig. 1d-f). These results suggest that pTrkB in axons of mature pyramidal neurons predominantly locates in signaling amphisomes which are absent in developing neurons, where TrkB is localized to signaling endosomes^22,23^.

We next asked whether synaptic activity has impact on amphisome biogenesis. We used a sustained physiological stimulation protocol of 900AP@10Hz that mobilizes the total recycling SV pool and maximizes presynaptic membrane dynamics^24, 25^. Under moderate frequencies of 10Hz hippocampal synapses are capable of sustain neurotransmitter release by maximizing presynaptic membrane dynamics without causing significant alterations in the exo/endocytosis rates^26^. We refer to this protocol as high-fidelity stimulation (HFS). In comparison to cultures silenced with TTX, we observed an increased number of axonal LC3 puncta 30 min following HFS in axons at DIV15 (Fig. 1k,l), indicating that this protocol induces robust synaptic autophagy. In differentiated neurons HFS increased amphisome number as evidenced by the overlap of LC3/pTrkB immunolabel while silencing of cultures with TTX had the opposite effect (Fig. 1m,n). Collectively these data show that amphisome biogenesis is tightly linked to synaptic activity.

### Autophagy and the endolysomal pathway are segregated in axons

Mature lysosomes are apparently present in axons of mature neurons^27^. Previous work suggests that amphisomes retain their signaling capacity during retrograde transport to the soma^20, 21^, which requires evasion from the fusion with axonal lysosomes. We indeed found very little co-localization of amphisomes with the lysosomal markers LAMP2A (Fig. 1o,p) and Cathepsin D (Fig. 1s-t). Moreover, no increase of lysosomal membranes was apparent in distal axons following HFS (Fig. 1q-r), and the number of autolysosomes detected by LC3 vesicles positive for LAMP2A or Cathepsin D in response to enhanced synaptic activity remained unchanged (Fig. 1r,u-v). Thus, amphisomes are devoid of degradative properties and the autophagy and endo-lysosomal pathway remain segregated in axons irrespective of synaptic activity.

### Phagophore formation is initiated at active presynaptic boutons

In *Drosophila* neuromuscular junctions autophagosome formation has been observed at active boutons^28^, however, the is no evidence for activity-dependent autophagy induction at presynaptic sites in mammalian neurons^11^. We reasoned that this might reflect the low abundance of autophagy-related proteins in the synaptic proteome^29^ and the transient nature of activity-dependent autophagosome formation at boutons. To address these issues, we employed a novel monoclonal antibody that specifically detects phosphorylated autophagy-related protein 16-like 1 (pATG16L1)^30^. ULK1 kinase directly phosphorylates and activates ATG16L1^31^, which as part of the autophagy-related ATG12-ATG5/ATG16 complex is responsible for coupling ATG5-ATG12 conjugate to membranes of nascent autophagosomes, and in turn for LC3 (ATG8) lipidation and autophagosome formation^32^. The ATG16L1/ATG5-ATG12 complex dissociates from the phagophore membrane before or around the time of autophagosome closure^32, 33^. Thus, pATG16L1 immunofluorescence level provides a reliable measurement of autophagy rates and directly reflects autophagic vesicle formation. With this tool at hand, we next looked at the spatio-temporal dynamics of activity-dependent autophagy formation at boutons. We found a transient up-regulation of presynaptic pATG16L1 immunofluorescence in response to HFS at 15 minutes (Fig. 2a-b). Pharmacological inhibition of ULK1/2 with SBI-0206965 prevented HFS-induced ATG16L1 phosphorylation (Fig. 2a-b). Stimulated Emission Depletion (STED) nanoscopy revealed that pATG16L1 is found in close association to the presynaptic scaffolding protein Bassoon (Fig. 2c) and not detectable in postsynaptic dendritic spines (Fig. 2d). Transmission electron microscopy (TEM) provided direct evidence for autophagy initiation at presynaptic terminals (Fig. 2e-f) and we identified an increased number of a variety of circular membrane structures resembling autophagy-related vesicles in presynapses following HFS (Fig. 2f). No change in pATG16L1 fluorescence was found at postsynaptic sites labeled by Shank3 (Extended Data Fig. 2a-c), altogether indicating that phagophore formation following HFS is restricted to presynaptic boutons. Along these lines we observed that pATG16L1 immunofluorescence intensity was clearly correlated with neurotransmitter release as revealed by a synaptotagmin-uptake assay^34^ (Extended Data Fig. 2d-f).

**Figure 2.**
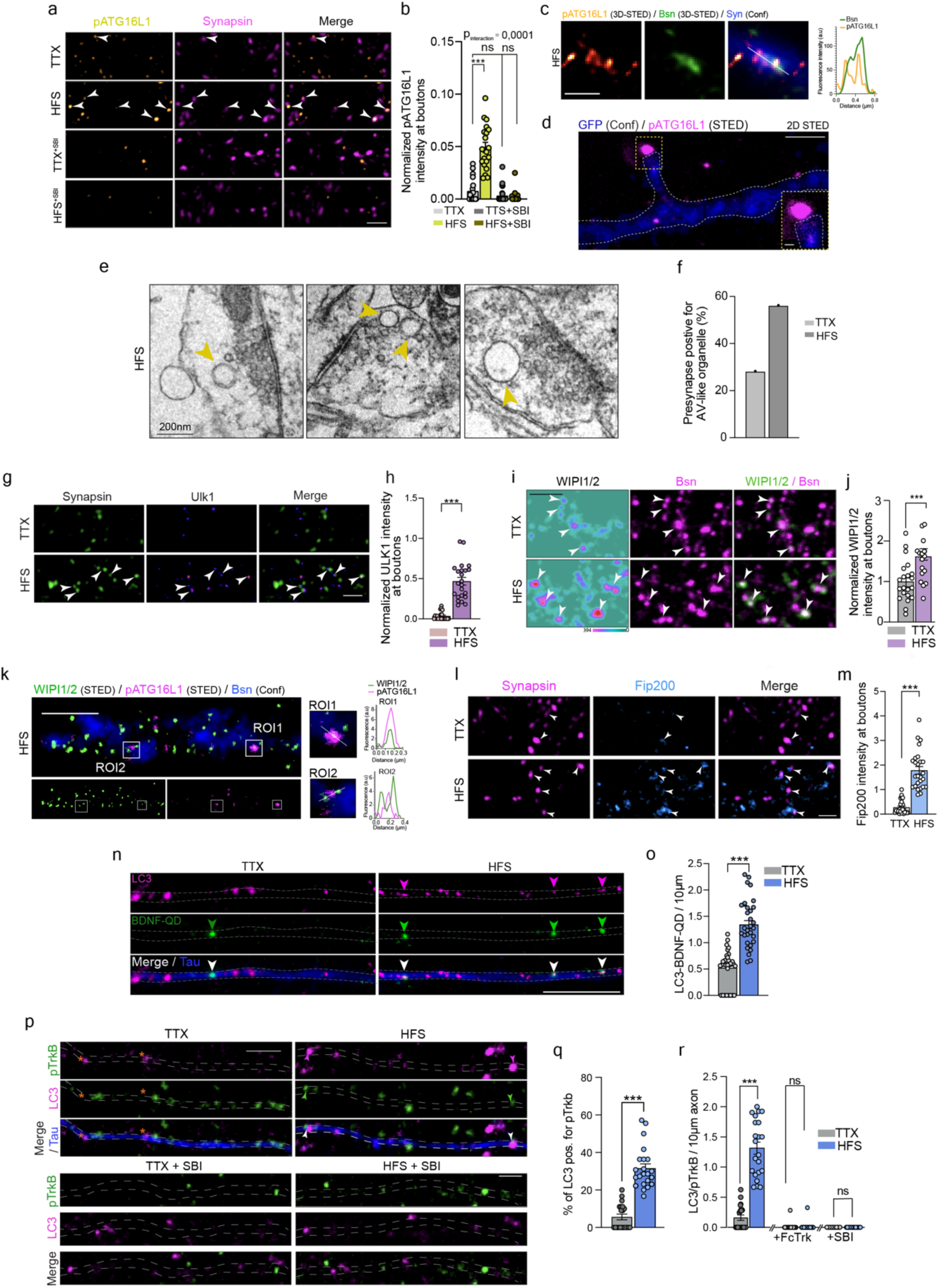
Electrical field stimulation causes autophagy activation at boutons. A) Representative images of TTX or HFS-treated primary neurons immunostained for Bassoon and pATG16L1. B) Bar graph depicting the intensity of pATG16L1 at the presynapse identified by Bassoon in neurons treated with TTX, HFS and HFS after 2h of treatment with SBI-0206965. (*n=* *20-27 from 3 independent cultures.) Statistical analysis was performed using two-way ANOVA followed by Tukey’s multiple-comparisons test*. C) Superresolution 3D STED image of pATG16L1 localizing with the presynaptic scaffold Bassoon. Synapsin signal acquired with confocal microscopy is shown in blue. Left, intensity line profile from the images. Scale bar is 500nm. D) 2D STED image displaying GFP as neuronal cell fill, and pATG16L1. Scale bar indicate 1µm and 200 nm (inset). E) Representative electron micrographs from synapses of HFS-treated primary cultures. F) Bar graph depicting the percentage of presynapses positive for AV-like organelles from 2 different experiments. G) Representative images of the presynapse labelled with Bassoon and Ulk1. Scale bar indicates 2µm. H) Quantification of Ulk1 intensity measured in Bassoon-positive boutons from TTX and HFS groups. *n=18 from 3 independent cultures* I) Representative image of primary neurons immunostained against WIPI1/2 at boutons labelled with presynaptic marker Bassoon. Scale bar indicates 2µm. J) Quantification of WIPI1/2 intensity at boutons at TTX-treated and HFS groups. *n=18 (FS) n=16 (TTX) from 3 independent cultures* K) 2D STED image from HFS cells stained for WIPI1/2 and pATG16L1, as well as bassoon in confocal. White squares in main images indicates ROI1 and ROI2 that are zoomed in on the right side accompanied by line profiles showing fluorescence intensity over distance. Scale bar indicates 1µm. L) Representative images showing immunostaining of primary neurons against FIP200 and Synapsin as presynaptic marker for TTX and HFS conditions. Scale bar indicates 2µm. M) Bar graph depicts FIP200 intensity measured at presynapses detected by Synapsin comparing TTX and HFS conditions. *n=21 (FS) n=23 (TTX) from 3 independent cultures* N) Representative images of neurons incubated with BDNF-QD655 and immunostained against LC3. Scale bar indicates 5 µm. O) Quantification of amphisome number in 10µm axon detected as LC3/BDNF-QD655. *n=30 from 3 independent cultures* P) Representative image of primary neurons treated with TTX or HFS and immunostained against LC3, pTrkB and Tau. Below, representative images of primary neurons treated with SBI-0206965 and immunostained against LC3, pTrkB and Tau. Scale bar indicates 2µm. Q) Quantification of the total percentage of LC3 puncta positive for pTrkB showing the relative amount of amphisomes to autophagosomes in 10 µm axon of TTX vs HFS cells. (*n=22 (FS) n=20 (TTX) from 3 independent cultures)* R) Number of colocalizing LC3/pTrkB puncta in 10 µm axon in TTX and HFS cells (*n=22 (FS) n=20 (TTX) from 3 independent cultures)* and quantification in cells treated with SBI-0206965 for 2h (*n=16 (FS) n=14 (TTX) from 2 independent cultures*) and Fc Trk overnight (*n=26 (FS) n=25 (TTX) from 3 independent cultures)*, respectively. Data are plotted as average ± SEM. ^∗∗∗^ represents p < 0.0001 to 0.001 by two-tailed Mann-Whitney U test.

### Core components of autophagy-machinery are recruited to presynaptic boutons in response to sustained synaptic activation

A substantial increase in the number of LC3-positive vesicles, likely representing mature autophagosomes in axons, is detected about 30 min following HFS (Fig. 1k,l) raising the question why there is a certain delay in phagosome maturation. We next assessed whether key components of the autophagy machinery are recruited directly to boutons after stimulation of primary cultures. Within 15 minutes the autophagy activating enzyme ULK1 is prominently recruited to the active zone (Fig. 2g,h). Similarly, but to a lesser extent WIPI1/2 proteins, that are required in mediating the PI3P signal at the onset of autophagy^32^, were more prominent at boutons (Fig. 2i,j) in close proximity to pATG16L1 as revealed by STED nanoscopy (Fig. 2k). In addition, immunofluoresence of the ULK1-interacting protein FIP200 (Fig. 2l,m), the lipid scramblase ATG9 (Extended Data Fig. 2g-i) and the autophagy receptor p62 (Extended Data Fig. 2j,k) was increased following HFS. In particular ULK1 and FIP200 are recruited rapidly, i.e. within minutes, to presynaptic boutons and phagophore formation may start very close to the active zone defined by the presence of the CAZ protein Bassoon (Fig. 2k).

### Signaling amphisomes are organelles of the canonical autophagy pathway

As expected, a much higher fraction of axonal LC3-positive vesicles contained BDNF-qDot655 (Fig. 2n,o) as well as active pTrkB following HFS (Fig. 2p,q; Extended Data Fig. 2l,m). When endogenous BDNF was chelated with Fc-bodies no amphisomes were detected in neurons silenced with TTX or after HFS (Fig. 2p,r; Extended Data Fig. 2n,o). LC3-associated endocytosis (LANDO) is a noncanonical function of the autophagy machinery, in which LC3 is conjugated to Rab5-positive endosomes, using a portion of the canonical autophagy pathway that is independent of ULK1 activation^35^. At variance to LANDO, however, presynaptic amphisome biogenesis could be blocked with the ULK1-inhibitor SBI-026965 (Fig. 2p,r; Extended Data Fig. 2p).

With correlative light electron microscopy (CLEM) we then confirm the localization of TrkB in double-membrane organelles suggesting macroautophagy is induced at boutons in an activity-dependent manner. Primary cells were fixed following HFS and confocal microscopy helped to identify vesicles positive for both, LC3-GFP and TrkB-RFP in axons (Fig. 3a). Several regions of interests were selected and ultrastructure analysis of those ROIs by means of electron microscopy revealed the presence of round organelles of ∼400-500nm in diameter with a heterogeneous content. Importantly, in all cases these organelles exhibited a double-membrane ultrastructure indicative for amphisomes of the canonical macroautophagy pathway and not the endolysosomal pathway (Fig. 3a-a’’).

**Figure 3.**
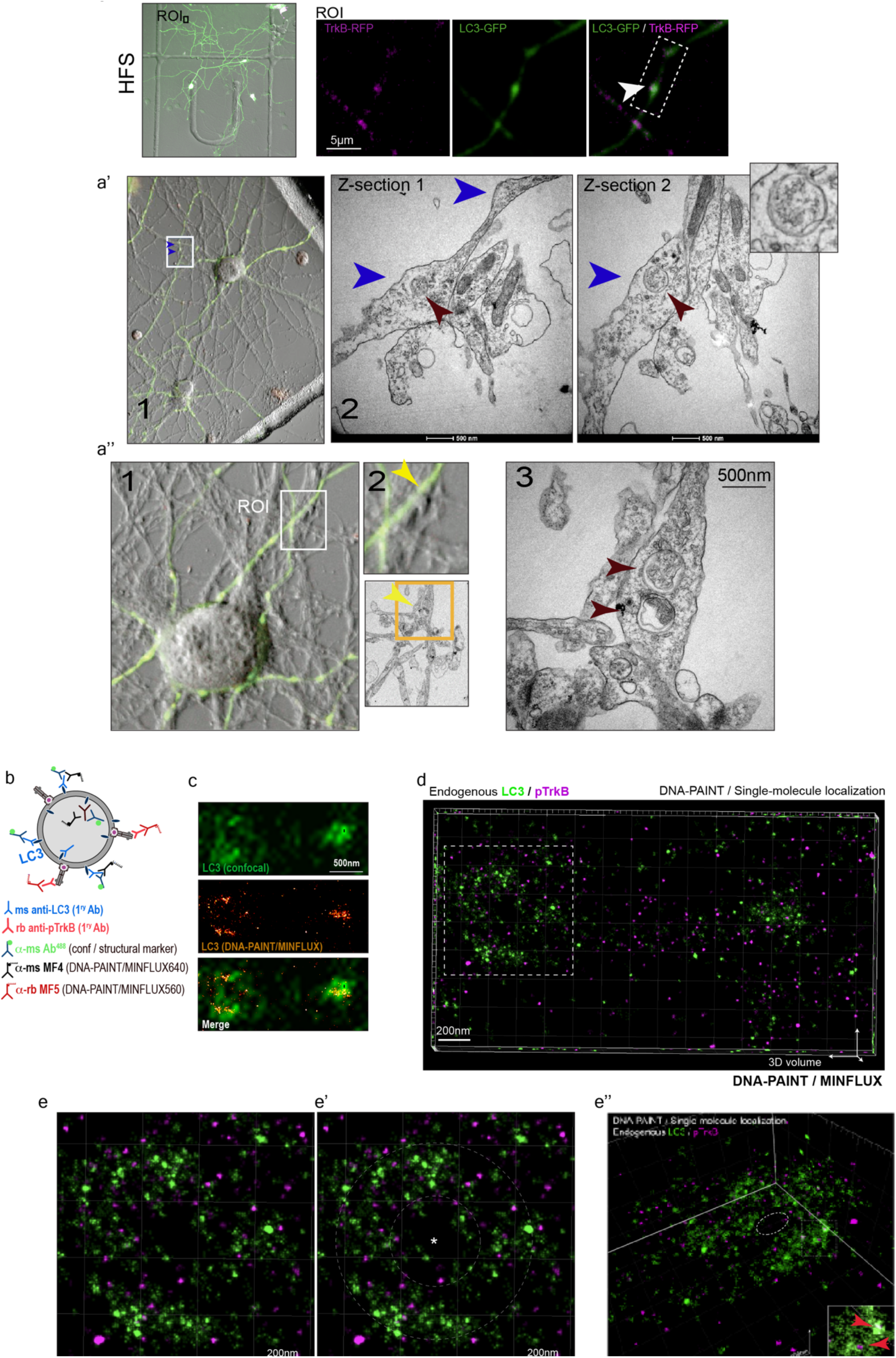
Ultrastructure of amphisomes revealed by DNA-PAINT/MINFLUX. A) Representative images of the experimental approach employed to visualized amphisomes using correlative light-electron microscopy (CLEM). The overview image in the left is a merge image acquired with transmitted light (TL) and fluorescence (LC3-GFP and TrkB-RFP) microscopy. Co-localizing LC3/TrkB puncta were selected as region of interest (ROI) by visual inspection of axons. A selected ROI is shown in the zoomed-in images on the right. **A’** shows an overview image of the regions were the ROI was selected (1). On the right side (2), electron micrographs of the selected ROI. Blue arrows indicate the same regions in confocal microscopy and electron microscopy. Please note that the electron micrographs are tilted to the right with regard to the confocal image. Two different Z-planes of the same region are shown as Z-section 1 and 2. Red arrow indicate an amphisome which is shown at a higher magnification in the upper right corner. **A’’** shows a second CLEM example. 1 shows an overview image of the selected ROI and the corresponding electon micrograph is shown in 2. Image nr 3 shows a higher-resolution electron micrographs where two amphisomes are indicated by red arrows. B) Schematic of the experimental approach used in the experiments. Two primary antibodies against LC3 and pTrkB were used in primary neurons stimulated by HFS. Two secondary antibodies against ms LC3 (one used as structural marker and the other one for DNA-PAINT) and one against rb pTrkB were used. C) Endogenous LC3 imaged by confocal (upper panel) and DNA-PAIN/MINFLUX. D) 3D volume of single-molecule localizations obtained for endogenous LC3 and pTrkB with DNA-PAINT/MINFLUX. Scale bar is 200nm. E) Zoomed image of LC3 and pTrkB 3D localization from the boxed ROI shown in D (scattered square). Scale bar is 200nm. In E’ the asterisk indicates the empty lumen and the dounought structure is indicated by the lines. In E’’ a 3D volume plot of single-molecule localizations obtained for LC3/pTrkB from the structure in E. The inset shows the tight association of pTrkB with LC3 as indicated by the red arrows. Pixel size is 4nm. Scale bar is 200nm.

To gather further structural information, we employed minimal photon fluxes (MINFLUX) nanoscopy^36^ to visualize amphisome ultrastructure in neurons. To this end we employed DNA-PAINT, a labelling technique used in single-molecule localization microscopy, to detect the localization and distribution of endogenous LC3 and pTrkB receptors. When combined with MINFLUX, DNA-PAINT reaches a localization precision of ∼3nm, allowing the visualization of structures at the molecular scale. Primary antibodies against LC3 were immunodetected by two secondary antibodies in parallel: αms-488, which provided a fluorescent signal in confocal microscopy and served as structural marker, and αms-MF4, which was employed for DNA-PAINT/MINFLUX acquisitions (Fig. 3b). LC3-positive structures were selected based on the signal acquired in confocal microscopy. Subsequent 3D-single-molecule localizations of LC3 in these structures were obtained using DNA-PAINT/MINFLUX imaging (Fig. 3d,c,e). A second acquisition using DNA-PAINT/MINFLUX was performed to obtain localizations of single pTrkB molecules in the same organelles. DNA-PAINT/MINFLUX recordings were performed for several hours to enhance the number of localizations per channel, and only localizations that had been repeatedly found in >4 scans for each channel were considered. We found LC3-positive structures with sizes ranging from 300-600nm, which indicates the existence of autophagy vesicles at different stages (Fig. 3e; Extended Data Fig. 3a-c). In larger vesicles, LC3 is organized as a halo of ∼80nm in thickness, resembling a doughnut with an inner space devoid of LC3 or pTrkB (Fig. 3e-e’’). Given that the thickness of the membrane lipid bilayer ranges from 5-10 nm and that the extension of antibodies is ∼15nm (i.e. ∼30nm for primary/secondary antibody complexes), the area occupied by LC3 in these organelles is consistent with double-membrane organelles, where LC3 is located in the inner and outer membrane and where the inner lumen contains degradative cargo (Fig. 3e-e’’ and Movie 1). Importantly, we found pTrkB closely associated with LC3, confirming previous findings^20^, as well as the double-membrane nature of TrkB-signaling amphisomes further indicating that amphisome biogenesis is independent of LANDO (Fig. 3e’’). Of note, even in smaller vesicles,

LC3 was prominently found occupying the organelle lumen (Extended Data Fig. 3b,c), supporting the presence of LC3 inside amphisomes. Taken together, this set of experiments shows that activity-dependent presynaptic autophagy results in the formation of predominantly signaling amphisomes.

### Bulk endoyctosis serves as membrane donor for amphisome formation

During high frequency stimulation SV endocytosis triggers activity-dependent bulk endocytosis (ADBE) to increase the retrieval capacity within the nerve terminal^37^. A quantitative assay of ADBE uses uptake of fluorescent dextrans of high molecular weight (40kDa) as fluid phase markers^38^. HFS induced bulk endocytosis of presynaptic membrane as evidenced by increased dextran fluorescence intensity at boutons (Extended Data Fig. 4a-b) and dextran puncta along axons (Fig. 4a-b). These puncta prominently colocalized with LC3 and pTrkB (Fig. 4a,c). Similarly, VAMP4, which is selectively recruited to large bulk endosomes^39^, showed increased co-localization with amphisomes identified by means as LC3/BDNF-qDot positive (Fig. 4d-f). We next assessed whether ADBE is a requirement for amphisome biogenesis by employing a TAT-peptide that, in a dominant-negative way, specifically prevents ADBE without affecting clathrin-dependent endocytosis^40, 41^. The peptide corresponds to the sequence of the dynamin I phosphobox, where two serine residues are replaced by alanine to bind syndapin (TAT-AA), while the phosphomimetic substitution (TAT-EE) has reportedly no effect^40^. Incubation of neurons with TAT-EE completely prevented HFS-induced dextran uptake at boutons, whereas TAT-AA did not affect dextran uptake following HFS (Fig. 4g,h). Most important, inhibition of ADBE induced by the TAT-EE peptide completely blocked amphisome formation in response to HFS (Fig. 4i,j) suggesting an essential role of bulk endocytosis for phagophore formation at boutons.

**Figure 4.**
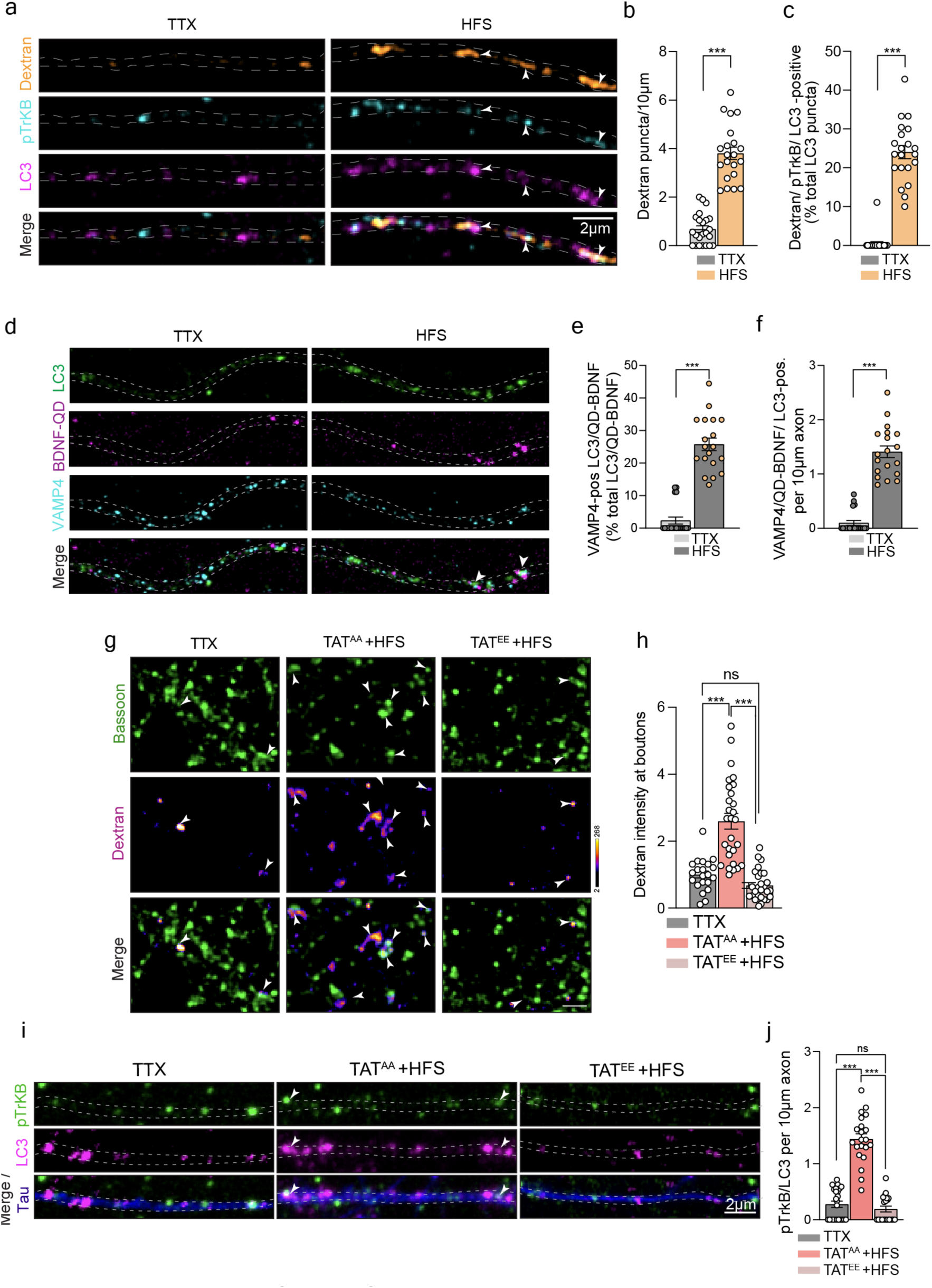
Amphisomes are formed from bulk-endosomes. A) Representative images of primary hippocampal neurons at DIV14 incubated with high-molecular weight Dextran (40kDa) that were either treated with TTX or stimulated by means of HFS in the presence of dextran. Stained for colocalization analysis with amphisome markers LC3 and pTrkB. Scale bar indicates 2µm. B) Bar graph shows number of Dextran puncta along 10µm of axon from A. C) Quantification of the total percentage of analyzed LC3 puncta along axons positive for Dextran in TTX treated to HF-stimulated cells. (B+C) *n=23 (HFS), n=22 (TTX) from 3 independent cultures* D) Representative image of neurons at DIV14 labelled with BDNF-QD655 and immunostained for LC3 and VAMP4 for TTX and HFS. E) Quantification of the total percentage of amphisomes labeled with BDNF-QD655 and LC3 and positive for VAMP4 relative to total LC3. F) Number of amphisomes positive for VAMP4 from D-E per 10µm of axons. *(E+F) n=20 from 3 independent cultures;* *(A-F)* Data are plotted as average ± SEM. ^∗∗∗^ represents p < 0.0001 to 0.001 by two-tailed Mann-Whitney U test. G) Representative image of primary neurons incubated with Dextran and stimulated in the presence of TAT-AA (mock) or TAT-EE and immunolabeled with the presynaptic marker Bassoon. Scale bar indicates 1µm. H) Dextran intensity measured at presynapses identified by Bassoon. Quantification to D. (n=28 from 3 independent cultures) I) Representative image of axons identified by Tau from primary hippocampal neurons at DIV14 immunostained against pTrkB and LC3. Showing TTX, TAT scrambled (AA) and TAT peptide (EE) treated conditions. J) Colocalization analysis of pTrkB and LC3 in 10µm axon. Quantification to I). *n=20 (HFS-TAT EE), n=21(TAT AA), n=24 (TTX) from 3 independent cultures* *(G-J)* Data are plotted as average ± SEM. Statistical analysis was performed using the Kruskal–Wallis test followed by Dunn’s multiple-comparisons test. *** represents *p* < 0.0001 to 0.001.

### Synaptic autophagy is initiated upon increased energy demand and requires AMPK activation

Two cellular signaling pathways are known to control autophagy initiation: AMPK (adenosine monophosphate-activated protein kinase) and mTOR (mammalian target of rapamycin) signaling. AMPK is the primary cellular energy sensor that is activated when the ADP/ATP or AMP/ATP ratio is increased^42^. Its activation induces autophagy by 1) inhibiting mTOR protein kinase complex and 2) directly phosphorylating and activating ULK1^17–19^. Synaptic activity imposes large energy demands that are met by local ATP synthesis through glycolysis and mitochondrial oxidative phosphorylation. In fact, 55% of ATP produced in neurons is used within boutons and axon terminals^43^. We indeed found that HFS causes AMPK activation as evidenced by a FRET-based sensor whose FRET efficiency was measured at boutons that were identified by Synaptophysin tagged with RFP (Sphy-RFP) (Fig. 5a-c). Live-imaging revealed a local activation of AMPK at boutons within minutes, a time course that corresponded to increased pATG16L1 immunofluorescence at boutons (see Fig. 2a,b). Most important, we could block autophagy induction at synapses (Fig. 5d,e), as well as amphisome formation in response to HFS with Compound C (Fig. 5f,g), a pharmacological inhibitor of AMPK. Treatment with Compound C prevented autophagy induction and LC3 puncta increase following HFS (Extended Data Fig. 5a), while levels of pTrkB puncta were not affected and slightly elevated following HFS (Extended Data Fig. 5b). Moreover, expression of a dominant-negative form of AMPK (Extended Data Fig. 5c) prevented the induction of autophagy at boutons by HFS (Fig. 5h,i) as well as the formation of amphisomes (Fig. 5j,k). Taken together, these results suggest that bulk endocytosis, recruitment of autophagy proteins (primarily ULK1 as well as FIP200) and local activation of AMPK converge to induce synaptic autophagy (Fig. 5l).

**Figure 5.**
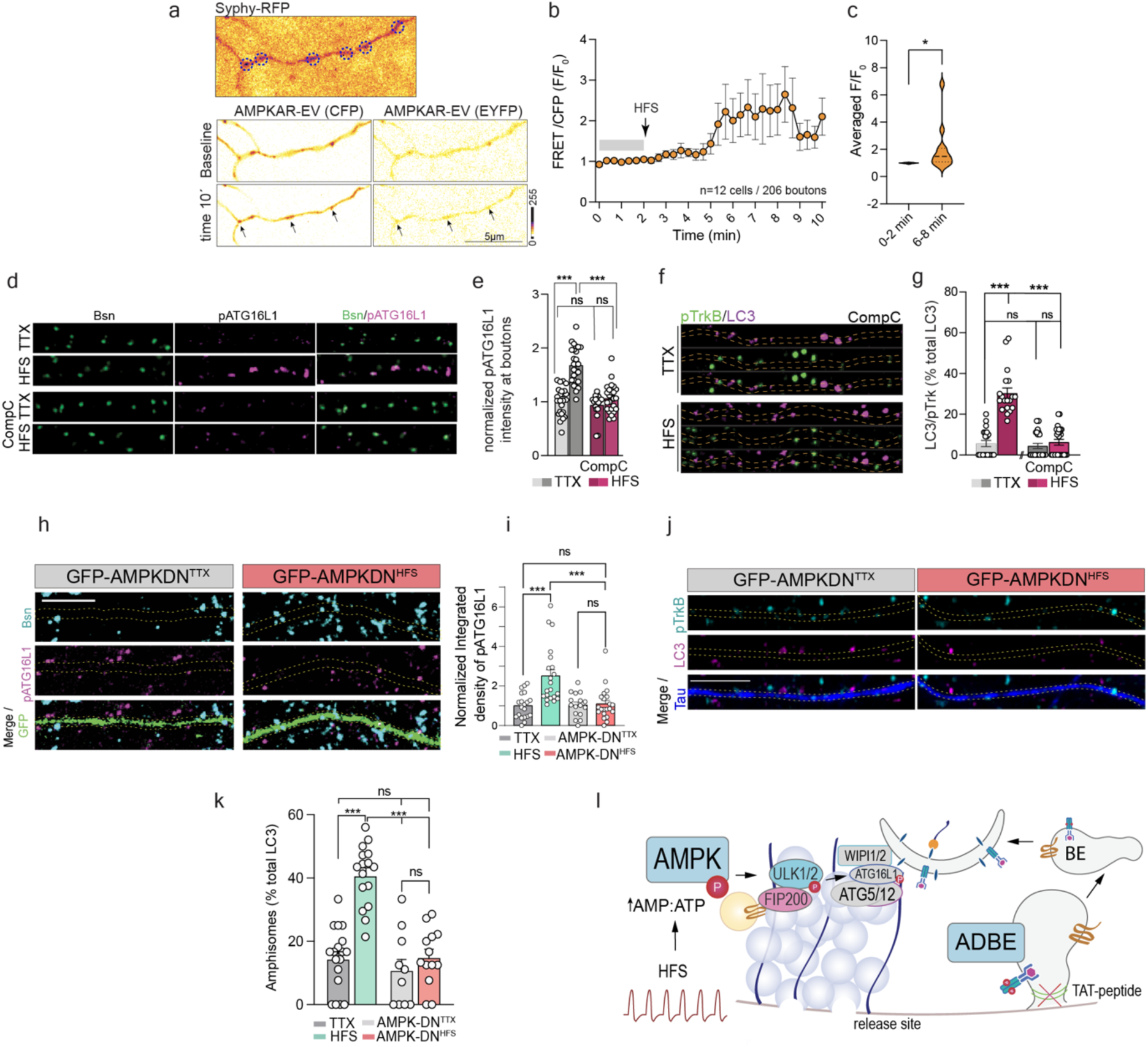
Synaptic autophagy and amphisome formation occur upon increased energy demands and AMPK activation. A) Representative images of primary neurons expressing AMPKAR-AV sensor at different time-points over the course of the experiment. Neurons co-expressed Sphy-tRFP together with AMPK-AV and Sphy-tRFP signal was to define presynaptic sites (dashed circles) where AMPK-AV fluorescence were measured (arrows). Scale bar is 5µm. B) Quantification of CFP/YFP as indicative of AMPK activation from the experiment in A. C) Averaged F/F0 values from the given time points in the experiment in A. Paired t-test was used as statistical analysis. D) Representative images of the primary neurons treated with TTX or HFS in the presence of the AMPK inhibitor Compound C (CompC) or DMSO as control and immunostained with Bassoon and pATG16L1. E) Quantification of pATG16L1 intensity measured in Bassoon-positive regions from the experiment in D *n=23(TTX); n=24(HFS); n=19(TTX+Comp.C); n=23(HFS+Comp.C) from 3 independent cultures* F) Representative images of of axons labeled by Tau from primary hippocampal neurons treated with CompC or DMSO at DIV14 immunostained against pTrkB and LC3 to identify amphisomes. G) Quantification of amphisomes as percentage of LC3-positive organelles in axons from the experiment in F. H) *(D-G) n=25(HFS+Comp C.); n=26 (TTX+Comp. C) from 3 independent cultures*Representative images of primary neurons expressing a dominant-negative form of AMPK tagged with GFP (GFP-AMPK-DN) and treated with TTX or HFS. After fixation, cells were immunostained for pATG16L1, Tau and Bassoon. I) Quantification of pATG16L1 intensity at presynaptic boutons labelled by Bassoon from the experiment in K. GFP-expressing cells were used as a control for comparison with the GFP-AMPK-DN group. J) Representative images of axons expressing GFP-AMPK-DN and immunostained against LC3/pTrkB following TTX or HFS for amphisome quantification. K) Quantification of amphisomes in GFP-AMPK-DN- positive or GFP-positive (control) axons from the experiment in M. L) Cartoon depicting the two processes underlying amphisome formation in the presynapse. Activity dependent bulk endocytosis and AMPK activation. (D-K) Data are presented as average ± SEM. Statistical analysis was performed Two-way ANOVA. ^∗∗∗^ represents p < 0.0001 to 0.001.

### Amphisome formation is linked to local *de novo* protein synthesis

Several recent studies demonstrate local translation in mouse presynaptic terminals at adult stages^2, 44–47^, detecting overall 1200 translated mRNAs *in vivo*^46^. Evidence has been provided that synthesis of proteins in presynaptic terminals may be involved in the local remodeling of the proteome in response to synaptic input, enabling a fast and specific regulation of the proteome in response to patterns of neuronal activity^48–51^. BDNF-TrkB promotes the release of mRNAs from RNA-binding proteins and these mRNAs become then readily available for translation^52, 53^. Moreover, RNA granules associate with late endosomes along axons and are sites of local protein synthesis^54^.

We therefore reasoned that amphisome biogenesis might couple synaptic autophagy to local protein synthesis and we indeed observed, employing a puromycilation assay, that de novo protein synthesis is increased locally at presynaptic sites in response to HFS (Fig. 6a,b). Interestingly, the puromycin label was associated with about 20% of axonal amphisomes (Fig. 6c,d), and HFS increased the number of autophagosomes that are in close proximity to ribosomes detected by immunolabeling of RPL10A (Fig. 6e,f) and RPS3A (Fig. 6g,h). This was further confirmed using STORM, which revealed an activity-dependent association of LC3-positive vesicles with ribosomes (Fig. 6i,j) and two-color STED nanoscopy (Fig. 6k).

**Figure 6.**
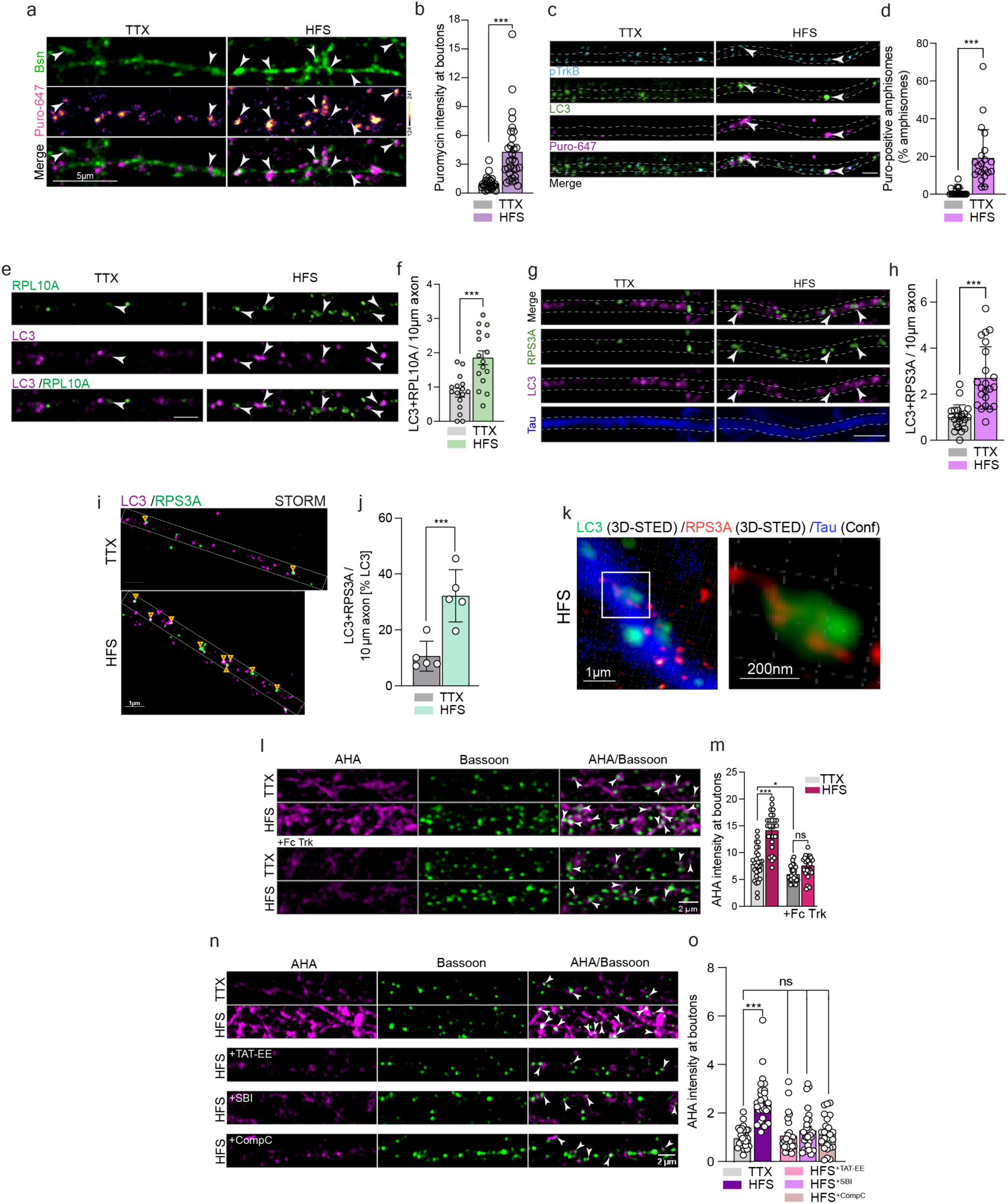
Amphisome formation is coupled to de-novo protein synthesis. A) Representative images of neurons treated with TTX or HFS in the presence of Puromycin-647 and immunostained against Bassoon. B) Bar graph depicting the intensity quantification of puromycin at the presynapse marked with Bassoon mask. *n=30(HFS); n=27 (TTX) from 3 independent cultures* C) Representative images of neurons treated with TTX or HFS in the presence of Puromycin-647 and immunostained against LC3 and pTrk for amphisome detection. D) Quantification of LC3 and pTrkB puncta positive for puromycin. *(n=21 (HFS); n=22(TTX) from 3 independent cultures* E) Representative images of neurons immunostained against LC3, RPL10A and Tau and treated with TTX or HFS. F) Quantification of the number of RPL10A-positive LC3 vesicles along axons from neurons treated with TTX or HFS. *(n=22 from 3 independent experiments.)* G) Representative images of neurons immunostained against LC3, RPS3A and Tau and treated with TTX or HFS. H) Quantification of the number of RPS3A-positive LC3 vesicles along axons from neurons treated with TTX or HFS. *(n=22 from 3 independent experiments.)* I) Representative images of axons from neurons treated with TTX or HFS and immunostained against RPS3A, LC3 and Tau. Samples were imaged with superresolution STORM nanoscopy. Image acquired with Abbelight SAFe MN360 in STORM using Abbelight Smart Kit buffer. Scale bar is 1µm. J) Quantification of the number of RPS3A-positive LC3 vesicles from the experiment in I. (A-J) Data are plotted as average ± SEM. ^∗∗∗^ represents p < 0.0001 to 0.001 by two-tailed Mann-Whitney U test. K) 3D-STED nanoscopy showing the close association of of RPS3A to LC3-positive structures in axons of HFS-treated neurons. L) Representative images of presynapses immunostained with Bassoon and newly synthesized proteins labelled with AHA over biotin-click reaction in neurons treated with TTX or HFS in the presence or absence of Fc-Trk bodies that chelate BDNF. M) Quantification of AHA intensity at boutons measured in Bassoon mask, comparing TTX condition to HFS of control and Fc bodies treated neurons from the experiment in K. *n=28 (TTX); n=27 (HFS); n=29(Fc TTX); n=26 (Fc FS) of 3 independent cultures*. N) Representative images of presynapses immunostained with Bassoon and newly synthesized proteins labelled with AHA over biotin-click reaction in neurons treated with TTX or HFS in the presence TAT-EE, SBI or CompC. O) Quantification of AHA intensity at boutons measured in Bassoon mask from the experiment in L. *n=29 (TTX); n=28 (HFS); n=27 (TAT); n=28 (SBI), n=27 (Comp.C) of* *3 independent cultures. (L-O)* Data are presented as average ± SEM. Statistical analysis was performed using the Kruskal–Wallis test followed by Dunn’s multiple-comparisons test. ^∗∗∗^ represents p < 0.0001 to 0.001, * shows p= 0.01 to 0.05.

FUNCAT labeling with the non-canonical amino acid AHA provided evidence that protein synthesis is increased at boutons when SV release was enhanced by HFS (Fig. 6l,m) and chelation of endogenous BDNF with Fc-bodies prevented this enhanced translation at boutons (Fig. 6l,m). Further evidence that biogenesis of BDNF/TrkB signaling amphisomes is instrumental for *de novo* protein synthesis at boutons was obtained from experiments showing that (i) suppression of bulk endocytosis by application of the TAT-EE peptide, (ii) inactivation ULK 1 with the antagonist SBI-026965, or (iii) blocking with AMPK activation using Comp C, suppressed HFS-induced enhancement of AHA-incorporation at presynaptic sites in hippocampal primary neurons (Fig. 6n,o).

### In *bsn-* and *sipa1l2*-deficient neurons synaptic autophagy is uncoupled from local protein synthesis

We next sought to determine the coupling of amphisome biogenesis to local protein translation in two loss-of-function genetic mouse models that interfere with different aspects of synaptic autophagy. The CAZ protein Bassoon reportedly inhibits synaptic autophagy by interacting with ATG5^55^. We indeed found much higher rates of autophagosome formation at boutons of Bsn-lacking neurons in TTX-silenced cultures (Fig. 7a-b) that remained at similar levels even after HFS (Fig. 7a-b). Similarly, the number of LC3-positive organelles in axons was increased in the absence of Bsn (Fig. 7c-d). This could reflect a ceiling effect preventing induction of additional activity-dependent autophagy. Accordingly, despite the reported higher levels of synaptic autophagy in the absence of Bsn^55^, we found that the number of BDNF/TrkB signaling amphisomes remained low and comparable to TTX-treated wild-type cells in *bsn*-deficient neurons (Fig. 7c-f). This finding further supports the need of bulk endocytosis and AMPK activation following sustained neurotransmission for amphisome biogenesis. We therefore used primary neurons from *bsn^KO^*mice to investigate whether amphisome biogenesis is a prerequisite for local activity-dependent protein translation. We were able to confirm this hypothesis by showing that AHA labeling does not increase at boutons in *bsn*-deficient neurons following HFS (Fig. 7g,h), reflecting the absence of activity-dependent amphisome formation. Following induction of presynaptic plasticity, amphisomes dissociate from dynein at boutons enabling local signaling and promoting transmitter release. This stop-over requires dissociation of SIPA1L2 from the dynein adaptor snapin^20^. Thus, although signaling amphisomes are formed in *sipa1l2*^-/-^ mice, they lose their capacity for local signaling at boutons^20^. To assess the impact of deficient local amphisome signaling on local protein synthesis, we first confirmed that autophagy is initiated at presynapses of *sipa1l2-/-* neurons following HFS (Fig. 7i,j). Next we confirmed that local protein synthesis depends upon local amphisome signaling as an increase in local protein synthesis was not detected in *sipa1l2*-deficient neurons following HFS (Fig. 7k,l). Moreover, re-expression of full-length SIPA1L2 tagged with GFP in *sipa1l2*^-/-^ neurons rescued this deficit (Fig. 7m,n). Thus, despite the activity-dependent formation of amphisomes, local signaling and probably docking to ribosomes, which is necessary to couple biogenesis of these hybrid organelles to activity-dependent protein synthesis at boutons, is interrupted in *sipa1l2*-ko neurons.

**Figure 7.**
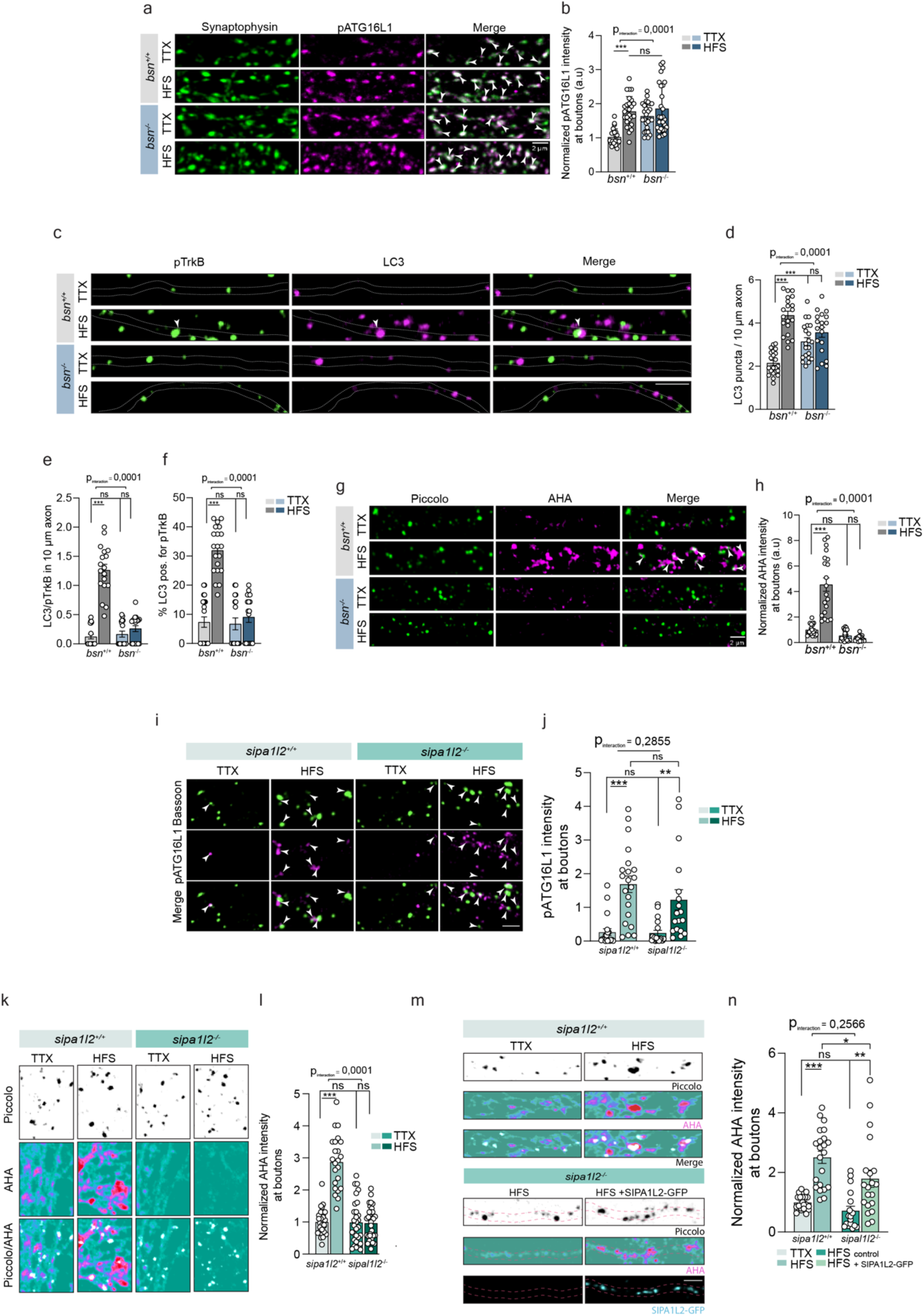
In *Bsn-* and *SIPA1L2*-deficient neurons activity-dependent amphisome formation is not coupled to local translation. A) Representative images of *bsn^+/+^* or *bsn^-/-^*neurons that underwent treatment with TTX or HFS immunostained against pATG1L1 and synaptophysin as presynaptic marker. B) Quantification of pATG16L1 intensity in synaptophysin-positive regions. *(n=24-27 from 3 independent cultures.)* C) Representative images of *bsn^+/+^* or *bsn^-/-^* neurons that underwent treatment with TTX or HFS immunostained against pTrkB, LC3 and Tau. D) Quantification of LC3-positive organelles per 10µm from the experiment in C. E) Quantification of amphisomes per 10µm axons from the experiment in A. F) Quantification of the percentage of LC3 organelles positive for pTrkB from the experiment in C (*n=24-27 from 3 independent cultures)*. G) Images represent presynapses immunostained with Piccolo and newly synthesized proteins labelled with AHA over biotin-click reaction from *bsn^+/+^* or *bsn^-/-^*treated with TTX or HFS. H) Quantification of AHA intensity at boutons measured in Piccolo mask, comparing TTX condition to high frequency stimulation of WT to KO mice. Bar graph to E. (*n=27-29 from 3 independent cultures)*. I) Representative images from *sipa1l2* wt and ko cells treated immunostained against Bassoon and pATG16L1 following TTX or HFS. Scale bar is 2µm. J) Quantification of pATG16L1 intensity in Bassoon-positive puncta from the experiment in I. K) Images represent presynapses immunostained with Piccolo and newly synthesized proteins labelled with AHA over biotin-click reaction from *sipa1l2^+/+^* or *sipa1l2^-/-^* treated with TTX or HFS. L) Quantification of AHA intensity at boutons measured in Piccolo mask, comparing TTX condition to high frequency stimulation of *sipa1l2^+/+^* or *sipa1l2^-/-^* mice. M) Images represent presynapses immunostained with Piccolo and newly synthesized proteins labelled with AHA over biotin-click reaction from *sipa1l2^+/+^* or *sipa1l2^-/-^* treated with TTX or HFS and expressing SIPA1L2-GFP. Neighbouring untrasfected cells were used as internal control. N) The graph shows the quantification of AHA intensity in SIPA1L2-GFP and neighbouring nontransfected cells from the experiment in M. *Data are presented as average ± SEM. Statistical analysis was performed using two-way ANOVA followed by Tukey’s multiple-comparisons test. pz values are indicated in the figure.* . ^∗∗∗^ represents p < 0.0001 to 0.001.

### BDNF-TrkB signaling amphisomes contain proteins of presynaptic nano-biomolecular condensates and drive translation of their corresponding mRNAs

In our TEM-analysis we barely found autophagosomes at presynapses that contained vesicular structures (Fig. 2e-f). We, therefore, asked about the potential degradative cargo in amphisomes. Multiple biomolecular condensates coexist at the presynapse to enable vesicle dynamics that underlie neurotransmitter release in the brain^56^. Synapsin-1 as well as Tau are known to form biomolecular condensates in the SV cluster that control the confinement and mobility of meso- and nanoscale SV domains, respectively^57, 58^. Components of the CAZ including Bassoon further coordinate the mobility and localization of recycling vesicles^56, 59^. Little is known about the local turnover of Synapsin, Tau and Bassoon among others despite playing a fundamental role in ensuring presynaptic structural and functional stability. Even though the presence of local mRNA in axons of all three proteins has been reported^2, 60^, their local translation has not been shown previously. Polyribosomes are scarce in axons and local translation of axonal mRNAs at boutons reportedly occurs mainly in monosomes^1^. Monosome-preferring transcripts often encode high-abundance synaptic proteins and preferentially mRNA of proteins that have a prominent presynaptic localization associate with ribosomes in axons^1^. Building on these previous results we first wondered which degradative cargo might be incorporated into amphisomes. Mass-spectrometry-based content profiling is for various reasons very difficult to achieve and outside the scope of a single manuscript. We nevertheless took a candidate approach and observed using STED nanoscopy that the cytoskeletal components of presynaptic biomolecular condensates Bassoon, Synapsin and Tau are present in LC3-positive vesicles (Fig. 8a-c) in axons following HFS.

**Figure 8.**
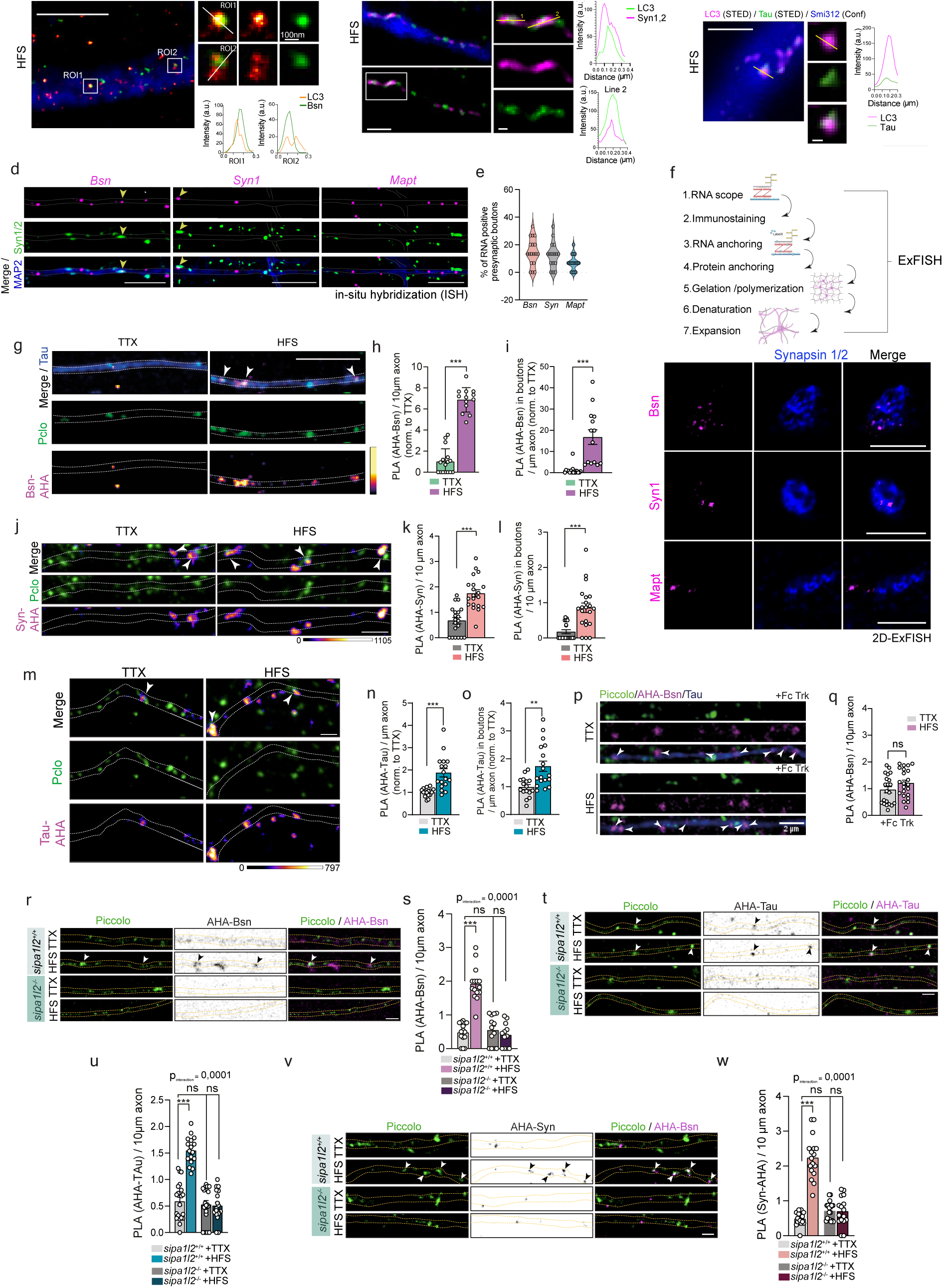
BDNF-TrkB amphisomes drive the degradation of key presynaptic proteins and the local translation of their mRNA. A-C) 3D-STED nanoscopy shows the colocalization of Bassoon (A; scale bar indicates 2µm and 100nm in inset), Synapsin (B; scale bar indicates 500nm and 100nm in inset) and Tau (C; scale bar indicates 500nm and 100nm in inset) and LC3 along axons detected with Tau (A-B) or Smi312 (C) in confocal resolution. D) Representative images of in-situ hybridization (ISH) experiments performed in primary neurons after HFS with probes targeting *Bsn, Syn1* and *Mapt*. Immunocytochemistry was performed afterwards for detection of synapsin and MAP2. Scale bar indicates 10µm. E) Quantification of the percentage of presynapses analyzed that contained the respective mRNA probles from the experiment in D (2 independent cultures). F) Explanatory scheme depicting the protocol used for Ex-FISH. Below, 2D ExFISH images displaying colocalization of *Bsn* and *Syn* mRNA probes with boutons labelled with Synapsin1/2 in HFS primary neurons. No colocalization of *Mapt* mRNA probe. Cortical presynaptic boutons in face (*Bsn*, *Syn*) and in side view (*Mapt*). Scale bar indicates 1µm. G) Representative confocal images of axons showing local translation of Bassoon by means of FUNCTA-PLA following HFS as compared to TTX-treated cells. Scale bar, 5 µm. White arrows indicate newly synthesized protein of interest. H) Quantification of PLA signal of locally synthesized Bassson per 10µm of axon from the experiment in G. I) Quantification of PLA signal in boutons labelled with Piccolo in axons per 10µm of axon from the experiment in G. *(H+I) n=22(TTX); n=19(HFS) from 3 independent cultures*. J) Representative confocal images of axons showing local translation of Synapsin by means of FUNCTA-PLA following HFS as compared to TTX-treated cells. Scale bar, 2 µm. White arrows indicate newly synthesized protein of interest. K) Quantification of PLA signal of locally synthesized Synapsin per 10µm of axon from the experiment in J. L) Quantification of PLA signal in boutons labelled with Piccolo in axons per 10µm of axon from the experiment in J. *(K+L) n=20(TTX); n=22(HFS) from 3 independent cultures*. M) Representative confocal images of axons showing local translation of Tau by means of FUNCTA-PLA following HFS as compared to TTX-treated cells. Scale bar, 2 µm. White arrows indicate newly synthesized protein of interest. N) Quantification of PLA signal of locally synthesized Tau in 10µm of axon from the experiment in M. O) Quantification of PLA signal in boutons labelled with Piccolo in axons per 10µm of axon from the experiment in M. *(N+O) n=22(TTX); n=19(HFS) from 3 independent cultures*. P) Representative confocal images and quantification of newly synthesized Bassoon at the presynapse labelled with piccolo in DIV14-16 cultured hippocampal neurons treated with Fc Trk bodies overnight. Q) Quantificaiton of the local translation of Bassoon upon HFS with Fc Trk bodies from the experiment in P. n=21(TTX); n=22(HFS) from 3 independent cultures *(G-Q) Data are presented as average ± SEM. ^∗∗∗^ represents p < 0.0001 to 0.001, ** shows p= 0.001 to 0.01 by two-tailed Mann-Whitney U test*. R-W) PLA signal of Bassoon (R,S), Tau (T,U) and Synapsin (V,W) of DIV18 cultured hippocampal neurons in *sipa1l2* WT and KO mice treated with TTX or HFS. *Sipa1l2^-/-^*neurons show no elevated PLA signal after HFS for newly synthetized Bassoon, Tau and Syn. (S) n = 17–19 cells from 2 independent cultures, (U) n= 17-20 from 3vindependent cultures, (W) n= 16-18 from 3 independent cultures. Scale bar indicate 2µm. Data are plotted as average ± SEM. *Statistical analysis was performed using two-way ANOVA followed by Tukey’s multiple-comparisons test. P values are indicated in the figure*.

We next wondered whether signaling amphisomes drive the translation of central components of biomolecular condensates. To address this question, we first confirmed the presence of the mRNAs of Bassoon, Synapsin and Tau at presynaptic sites. To this end we performed in-situ hybridization assays to *Bsn*, *Syn* and *Mapt* mRNA probes. Confocal microscopy revealed the presence of all mRNA at distal neurites and, to a lower extent, at presynaptic sites immunostained with Synapsin1/2 (Fig. 8d,e). To further confirm the presence of the candidate mRNAs in presynaptic sites we took advantage of expansion microscopy in combination with fluorescence in-situ hybridization (Ex-FISH) which allowed us to undoubtedly detect the presence of *Bsn* and *Syn1* mRNAs in presynapses (Fig. 8f). Transcripts of Tau (*Mapt*) were also found to be in very close association to presynaptic sites (Fig. 8f). We next established a proximity ligation assay (PLA) that allowed visualization of ongoing translation of these specific mRNAs at boutons (see Extended Data Fig. 6a). A clearly increased PLA signal in response to HFS was observed, indicating enhanced activity-induced translation of Bassoon (Fig. 8g-i; Extended Data Fig. 6b), Synapsin-1 (Fig. 8j-l; Extended Data Fig. 6c) and Tau (Fig. 8m-o; Extended Data Fig. 6d) at boutons labeled by immunostaining of the scaffold protein Piccolo, but also in axons labeled by Tau (Extended Data Fig. 6b-d). Importantly, no AHA-labeling and PLA signal was observed for transmembrane proteins, like the early endosome and lysosome marker LAMP1 or the cell adhesion molecule NCAM neither in TTX conditions nor after HFS (Extended Data Fig. 6e,f). Scavenging of endogenous BDNF with FC-bodies prevented enhanced local translation of Bassoon following HFS (Fig. 8p,q), indicating the involvement of BDNF/TrkB signaling in HFS-driven protein synthesis at boutons. Direct evidence that signaling amphisomes at boutons are crucial for activity-dependent translation of cytoskeletal protein components of the SV cluster was provided by experiments with *sipa1l2*-/- neurons. Gene knockout of this RapGAP resulted in a complete loss of activity-dependent de novo protein synthesis at boutons of Bassoon (Fig. 8r,s), Tau (Fig. 8t,u) and Synapsin (Fig. 8v,w).

## Discussion

One central question in the study of neuronal autophagy concerns the distinction between ongoing basal autophagy and autophagy induced by synaptic activity. In this study we have addressed this central question and our key findings reveal that i) biogenesis of signaling amphisomes indeed occurs at presynaptic boutons in an activity-dependent manner, ii) synaptic autophagy in hippocampal pyramidal neurons predominantly includes amphisome formation, iii) autophagic and endolysosomal pathways appear to be segregated in axons, iv) recruitment of proteins essential for autophagy initiation, bulk endocytosis and AMPK energy sensing converge for presynaptic phagophore formation, v) signaling amphisomes contain proteins of presynaptic nano-biomolecular condensates and drive local translation of their corresponding mRNAs (see Fig. 9).

**Figure 9:**
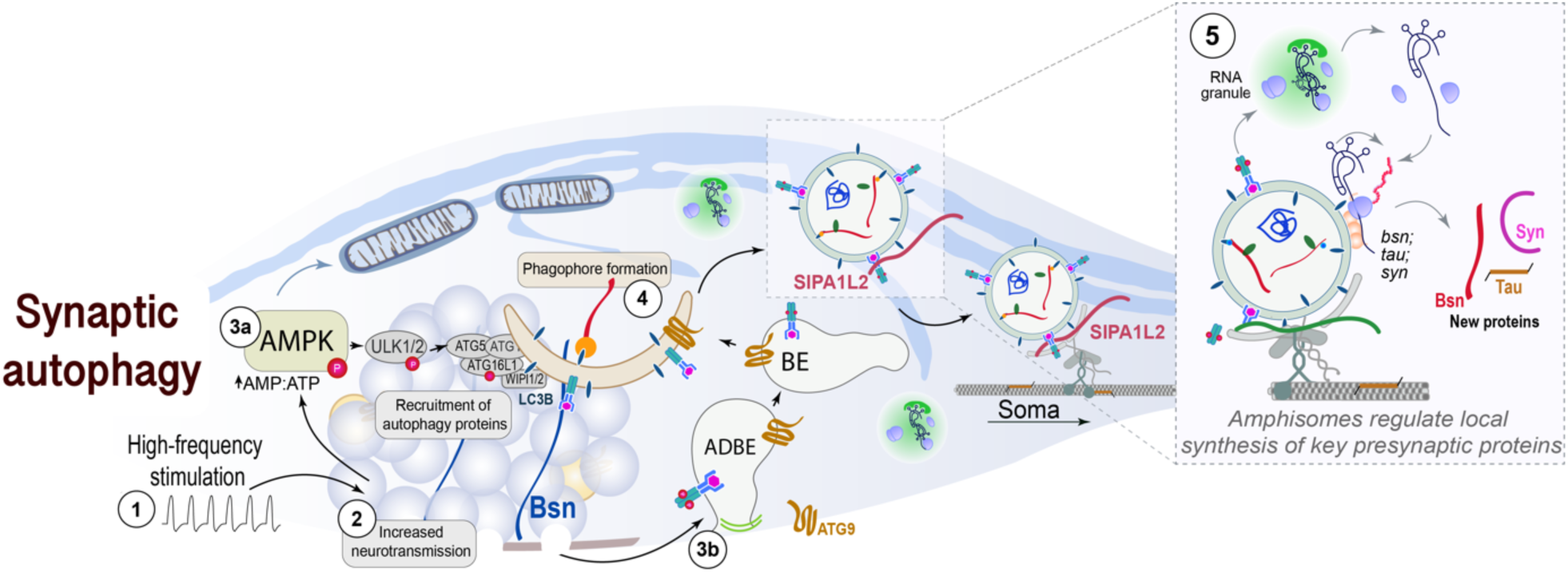
Underlying mechanism of amphisome formation at boutons. Under basal conditions smooth endoplasmic reticulum in axons and boutons is the most prominent site of continuous autophagosome formation. Following HFS (1), intense SV recycling (2) causes an increase in the AMP:ATP ratio that leads to the activation of AMPK (3a) and a recruitment of autophagy proteins to the bouton. AMPK triggers autophagy initiation at boutons. Concomitantly bulk-endocytosis takes place following HFS (3b) and the formation of a bulk endosome (BE) supports amphisome formation (4). BDNF/TrkB signaling from signaling amphisomes initiates the local synthesis of presynaptic proteins (5) essential for the maintenance and stability of presynaptic biomolecular condensates (including Bassoon, Tau and Synapsin) which, in turn, are likely to serve as cargo in amphisomes. In this way, signaling amphisomes enable the local replacement of essential presynaptic proteins.

### Induction of autophagy at boutons results in the formation of signaling amphisomes

Autophagy is widely seen as a self-degradative process that is important for balancing sources of energy at critical times in development and in response to nutrient stress. In neurons the situation is vastly different and it is evident that autophagy has to be under tight control in a compartment-dependent manner. Autophagy has the potential to engulf cargo to an extent that is detrimental for synaptic function. Therefore, cargo availability for engulfment has to be coupled to the mechanisms of autophagy induction, to ensure that the time and location of autophagosome biogenesis does not result in structural damage to boutons^11^. The convergence of three processes for induction of phagophore formation and highly localized engulfment of cargo led us speculate about a specific adaptation of autophagy to replenish the protein meshwork that configures SV cluster.

To study these processes for autophagy induction, we have used a physiologically relevant protocol for HFS that induces sustained reliable SV release and bulk endocytosis^24–26^. We suppose that bulk endosomes containing ATG9 and BDNF/TrkB receptors are formed upon HFS. After closure, a bulk endosome could fuse to an existing autophagosome to form a signaling amphisome. Alternatively, a bulk endosome might move closer to the ER at the bouton where ATG2 would mediate the transfer of phospholipids for autophagosome formation. In this scenario, the membrane content of the bulk endosome will become part of the signaling amphisome. A third scenario would be that bulk endosomes serve as a lipid source for an ATG9-containing carrier to grow into an autophagosome which will contribute to amphisome biogenesis.

### Mechanistic underpinnings of amphisome biogenesis

Our findings support the notion that synaptic autophagosomes are different from those generated in other neuronal sub-compartments, not only in terms of content, but also in terms of function. Content profiling will be necessary to provide a final answer whether selective or non-selective autophagy of degradative cargo is predominant at boutons. Interestingly, however, we barely found synaptic vesicles in autophagosomes at boutons and the data obtained suggest that the content of activity-induced synaptic autophagosomes is distinct from bulk autophagy under basal conditions that largely resembles ER-phagy^15, 61^.

Evidence for separate mechanisms underlying basal and activity-regulated autophagic processes in the presynapse comes from work on the *Drosophila* neuromuscular junction, where the regulation of activity-dependent degradation of SV components, but not of starvation-dependent autophagy, involves the vesicular membrane protein synaptogyrin^62^. The scarcity of autophagy-related proteins in the synaptic proteome^29^ and the lack of compelling evidence that autophagosomes are indeed formed in an activity-dependent manner at boutons in mammalia requested a systematic analysis with higher temporal and spatial resolution. Our results suggest that the phagophore is closed and cargo loading is completed at boutons within several minutes. This time course might at least in part reflect the necessity to recruit autophagy-related proteins to the presynapse. At present, it is unclear how sustained synaptic activity upon HFS relates to recruitment of ULK1 and FIP200 in particular, but one possible candidate is AMPK-signaling. We found that energy-sensing by AMPK is instrumental for phagophore formation and a pressing question concerns other processes that converge at boutons to generate amphisomes. Of particular interest is here the tight link between autophagic and endocytic factors^63–65^. Endophilin-A for instance, which is crucially involved in retrieval of membranes following the fusion SVs with the presynaptic active zone^66–68^ is also a major player in generating initiation sites for autophagy^28, 69^. The lipid phosphatase Synaptojanin-1 is recruited to Endophilin-A complexes and regulates the association of the lipid-binding protein ATG18a on nascent autophagosomes^70^. The AP-2 complex is involved in endocytosis during SV recycling^71, 72^ and can directly interact with ATG9A^73^, a lipid scramblase that is involved in early steps of phagophore formation^74^. Trafficking of ATG9 to boutons and insertion in the presynaptic membrane is a prerequisite for autophagy initiation and progression^75^. It will be interesting to study how these molecular interactions are related to bulk endocytosis, AMPK signaling and amphisome biogenesis.

### Amphisomes regulate mRNA translation of proteins of the SV cluster at boutons

A central finding of this study is that amphisome biogenesis connects synaptic autophagy to local protein synthesis. We propose that signaling amphisomes might couple energy demand due to increased SV release to rejuvenate the presynaptic protein pool. Sustained vesicular release likely comes along with increased protein damage and the necessity to replace proteins of the SV cluster. The low basal content of mRNA and protein synthesis machinery in axons depends on a rather local and efficient mechanism to replenish presynaptic proteins^76^. Signaling amphisomes could provide an efficient means to provide input specificity. Local translation offers a source for proteins that are used as close as 1–20 µm from their site of synthesis to elicit an effect^51, 76, 77^. The stop-over of amphisomes at boutons requires presynaptic activity and BDNF/TrkB signaling is restricted to highly active presynapses^20^. Moreover, signaling amphisomes appear to be highly stable organelles since the endolysosomal and autophagy pathway remain largely segregated in axons.

The mTOR pathway acts as a central regulator for metabolism by balancing protein synthesis and degradation. When mTOR is active it promotes protein synthesis while simultaneously suppressing autophagy^78^. Conversely, mTOR inhibition leads to the activation of autophagy. How can the same pathway at the same time activated to promote protein synthesis, be inhibited to induce autophagy? We suppose that there is a spatial segregation and that mTORC1 is absent from autophagy initiation sites at the active zone. AMPK can activate ULK1 directly and thereby trigger autophagy induction^78, 79^. Moreover, we were not able to induce autophagy in axons with rapamycin at a time scale relevant for this study (data not shown), supporting the idea that synaptic autophagy is not under control of the mTOR pathway.

### A role of Bassoon in activity-dependent synaptic autophagy, amphisome formation and local protein synthesis

Bassoon is the only currently known protein component of the CAZ that appears to regulate autophagy via an interaction with ATG5. However, apparent discrepancies exist in phenotypes of *atg5*- and *bsn*-deficient mice^15, 55^ and Bassoon is crucially involved in controlling presynaptic protein turnover in multiple ways that might indirectly interfere with the study of synaptic autophagy and that might accumulate over time. Lack of Bassoon increases BDNF/TrkB levels^80, 81^ as well as autophagy^55, 82^ and proteasomal activity^83^. We found that phagophore formation as evidenced by ATG16L1 phosphorylation is prominently enhanced at boutons of *bsn*-deficient cells in the presence of TTX and that HFS has no further effect on this measure as well as the number LC3-positive autophagic vesicles. We suppose that Bassoon controls activity-dependent synaptic autophagy and that therefore no further autophagosome formation can occur upon stimulation. In other terms, the inhibition of ATG5 function by Bassoon at rest seems to prevent elevated phagophore formation unless activity-dependent synaptic autophagy kicks-in. This, however, is a prerequisite for amphisome biogenesis in response to HFS.

### Functional consequences of amphisome signaling for the integrity of the SV cluster and nanodomains in biomolecular condensates

Local protein translation at boutons largely occurs at monosomes and despite some ribosome heterogeneity the emerging picture suggests that mainly highly abundant non-transmembrane synaptic proteins undergo activity-dependent de novo synthesis^1^. Interestingly, very few transmembrane proteins appear in the list of translated proteins at presynaptic monosomes^1^. Our results with a candidate approach support this view. The structure and integrity of the SV cluster crucially depend on cytoskeletal proteins, like Synapsin, Tau and proteins of the CAZ. These proteins form a dense meshwork that contains RNA^3^ and they undergo liquid-liquid phase separation to generate presynaptic nanoclusters whose density and number are regulated by activity. It has been proposed that they organize the nanoscale organization of SVs, a process that is still poorly understood. It is tempting to speculate that signaling amphisomes replenish proteins forming biomolecular condensates in a highly localized manner.

## Supplemental information

Supplemental information can be found online …

## Acknowledgments

The authors wish to thank Corinna Borutzki, Isabel Herbert, Monika Marunde and Peggy Patella for excellent technical assistance. MINFLUX nanoscopy was possible due to European Regional Development Fund (ERDF) under the Operational Programme Hamburg ERDF 2014-2020, REACT-EU, awarded by the Hamburgische Investitions- und Förderbank (IFB). Grant no. 51164232.

The authors received funding from the German Research Foundation (DFG) FOR5228 (RP06 to A.K. and M.R.K. Project number: 447288260 and RP03 to E.D.G.) and by ERC Starting Grant (LEXSYN) Number 101161748 (K.M.G.).

## Author contributions

M.A.A. and C.S. contributed equally to this work. M.A.A., A.V.F, T.M.B., E.D.G. and M.R.K. designed the experiments. M.A.A., C.S., C.M.V., S.M.W., C.S., R.T., K.M.G., R.B.A., L.M., S.Y., R.R., A.K., A.V.F. performed the experiments and analyzed the data. M.A.A. and C.S. prepared the figures. M.R.K. conceptualized the study and wrote the manuscript together with M.A.A. and E.D.G.. All authors read and commented on the manuscript.

## Material and Methods

A list of resources used in this study including antibodies, commercially available kits, chemicals and software, among other, is available in Table 1.

**Table 1.**
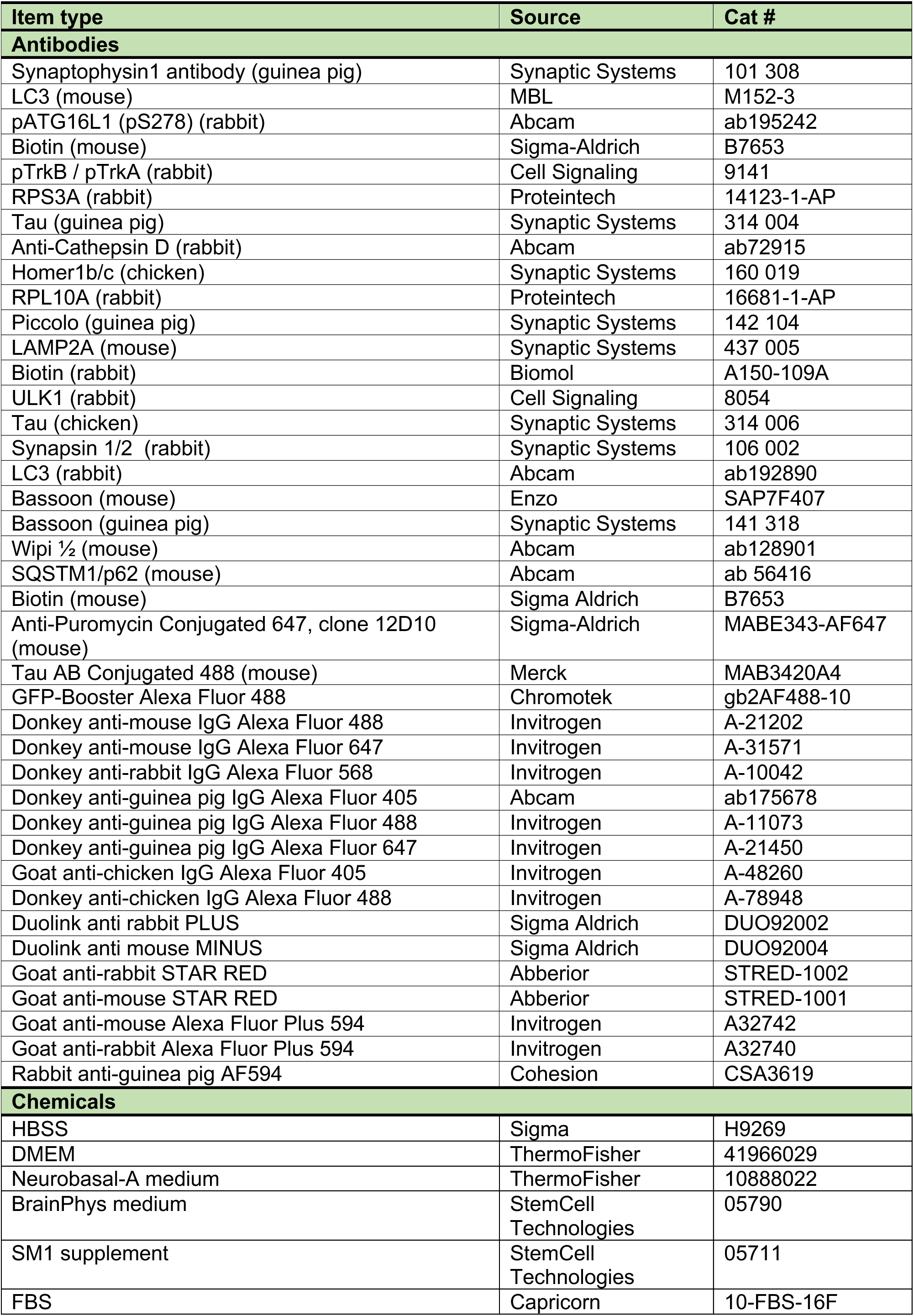
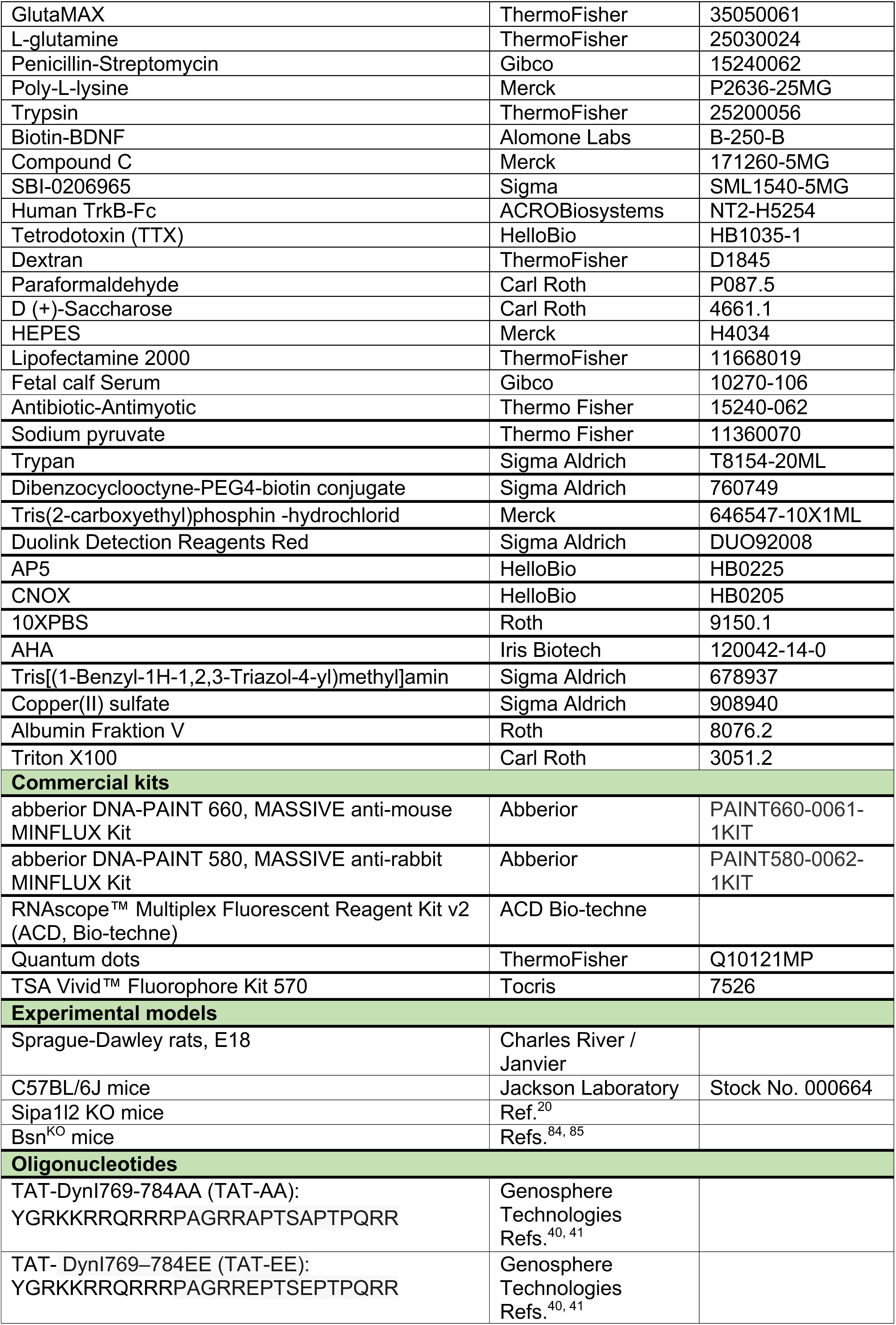
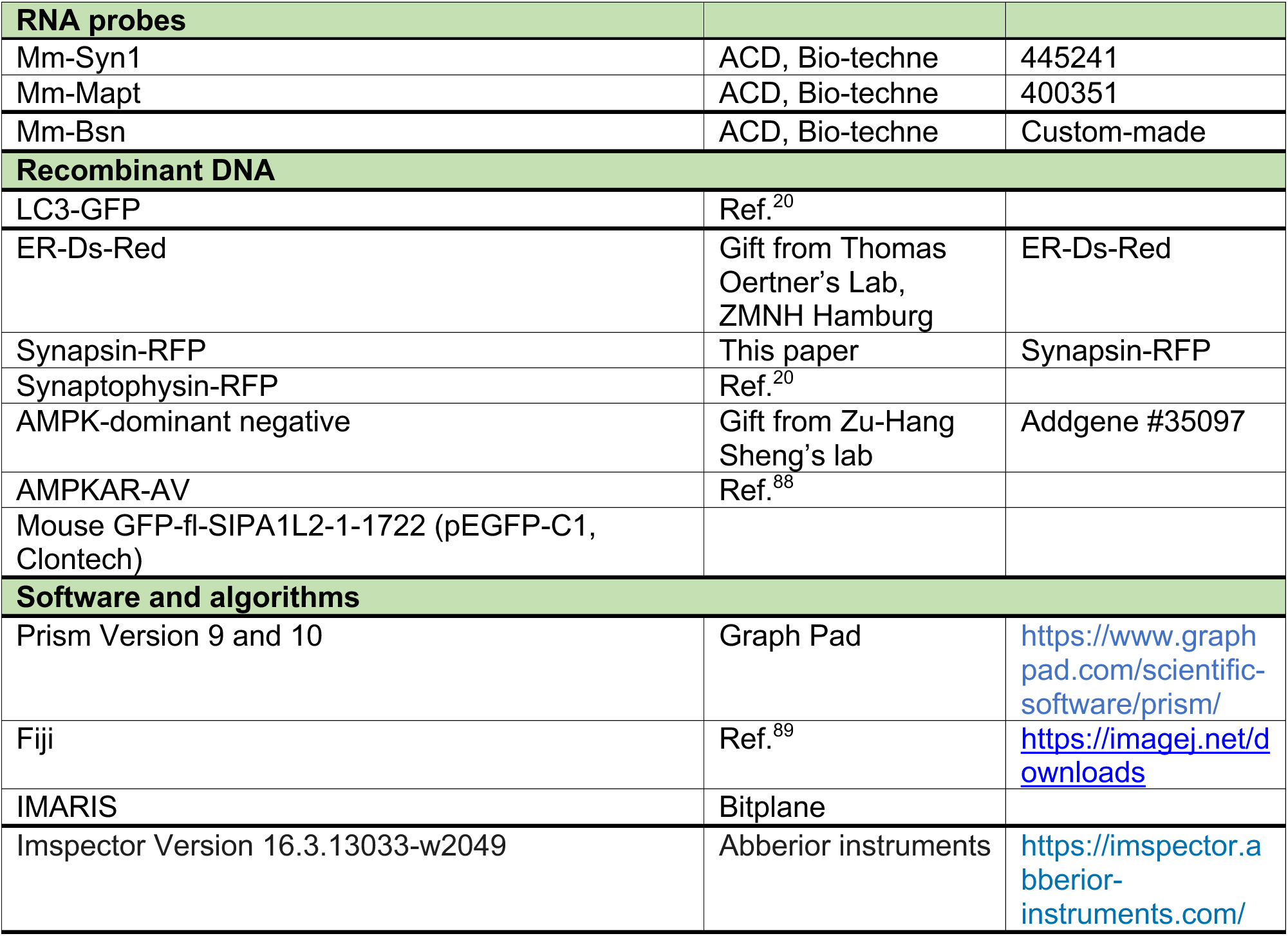
List of reagents, commercially available kits and software used in this study.

### Animals

Rats were maintained in the animal facility of the Leibniz Institute for Neurobiology, Magdeburg (Germany) or ZMNH, Hamburg (Germany) under controlled environmental conditions.

All animal experiments involving rats have complied with all ethical regulations for animal testing and research in accordance with the European Communities Council Directive (2010/63/EU) and were approved by the ethics committees of Sachsen-Anhalt/Germany (reference number 42502-2-1264 LIN and 42502-2-1284 UNIMD) or of the city-state of Hamburg (Behörde für Gesundheit und Verbraucherschutz, Fachbereich Veterinärwesen) and the animal care committee of the University Medical Center Hamburg-Eppendorf.

For experiments involving mice, breeding of animals and experiments using animal material were carried out in accordance with the European Communities Council Directive (2010/63/EU) and approved by the authorities of the State of Saxony-Anhalt (Landesverwaltungsamt Halle) to LIN (reference number: 42502-2-1797 LIN).

### Primary hippocampal cultures

All animal procedures complied with the ARRIVE guidelines, adhered to EU regulations, and were approved by the relevant institutional ethics committees. Sprague–Dawley rats were maintained in the animal facility of the Universitätsklinikum Hamburg-Eppendorf under standard housing conditions with unrestricted access to food and water. Experimental protocols were authorized by the local authorities of the State of Hamburg (Org 886; Nr. 125/17).

Primary hippocampal neurons were isolated from E18–E19 rat embryos of both sexes. Following decapitation, hippocampi were dissected in ice-cold HBSS and rinsed five times in the same solution. Tissues were then digested with trypsin at 37°C for 15 min, washed repeatedly with pre-warmed complete medium [DMEM supplemented with 1% L-glutamine, 1% penicillin–streptomycin, and 10% FBS], and subsequently dissociated mechanically using a 0.90 × 40 mm needle followed by a 0.45 × 25 mm needle . The resulting cell suspension was filtered through a 100 µm strainer, counted, and diluted in complete medium.

Cells were seeded at a density of 60,000 per 18 mm poly-L-lysine–coated coverslip and incubated at 37°C, 5% CO₂. After 1 h, the plating medium was replaced with BrainPhys supplemented with SM1 and L-glutamine. Neuronal cultures were maintained under these conditions, fed at DIV7 with an additional 200 µL of supplemented BrainPhys, and used for experiments between DIV14–16.

When required, preparation of hippocampal cultures from neonatal (P0-P1) mice was carried out. Sipa1/2^-/-^, Bsn^KO^, and their respective WT littermates were sacrificed by decapitation. Their brains were removed, and hippocampi were freed of meninges, plated one pair per 1.5 ml of Hibernate-A medium supplemented with 0.5 mM GlutaMAX™ and 2% B27 in a 2 ml Eppendorf tube, and stored at 4°C until PCR identified WT and KO animals.

Hippocampal tissue was treated with a final concentration of 0.25% trypsin, and the cell suspension was obtained by mechanical trituration. Cells were plated in DMEM with 10% FCS (fetal calf serum), 1mM glutamine, and antibiotics (100U/ml penicillin,100 ug/ml streptomycin at densities of 70.000-80.000 cells per coverslip (18 mm diameter). One hour after plating, the media were changed to Neurobasal A medium supplemented with 2% (v/v) B27, 1 mM sodium pyruvate, 4 mM GlutaMAX™, and antibiotics (100 U/ml penicillin, 100 μg/ml streptomycin). All neurons were maintained at 37°C in a humidified incubator containing 5% CO2. Cells were used for experiments between 14-16 DIV.

No formal statistical power calculation was performed, as sample sizes were not determined based on a predefined effect size. Instead, multiple independent experiments were conducted, each including the number of biological replicates indicated in the corresponding figure legends.

### Generation of knockout animals

Sipa1/2^-/-^ mice were generated as previously described^20^.

To generate constitutive Bsn^KO^ (Bsn^tm2Linmd^) mice, Bsn2^lx/lx^ (Bsn^tm1Linmd^) mice were crossed with a strain expressing ubiquitously Cre controlled by the CMV promoter, which resulted in Cre-mediated recombination in the germ line. Deletion of the exon 2 of the *Bsn* gene disrupts the reading frame of all exons downstream of exon 1, generating a premature stop codon and loss of the *Bsn* gene expression^84, 85^. To obtain the respective Bsn^KO^ and WT animals for our studies, heterozygous animals were backcrossed for 21-22 generations to the C57BL/6N background and then bred in a 2:1 ratio. For primary neuronal culture preparations and all other experiments, animals of both sexes were used.

### Plasmid DNA purification

Mini DNA purification was carried out by the alkaline lysis method. Overnight cultured bacteria were centrifuged for 5 min at 1000 x g and the pellet was resuspended in P1 buffer by vortexing. After the addition of P2 buffer, the tubes were incubated for 5 min at RT. P3 buffer was then added and after another 10 min incubation step at RT, the tubes were centrifuged at 16000 x g for 5 min at 4°C. 500 μl of the supernatant were transferred to isopropanol, incubated at RT for 10 min and centrifuged again as before. The pellet was washed with 70 % ethanol, centrifuged and then air-dried before solving the DNA in 30 μl Tris-HCl (pH 8.0). For large-scale DNA extraction, the NucleoBond Xtra Midi EF kit was used according to the manufacturer’s instructions.

### Transfections

Transfections were carried out simultaneously for both mouse and rat primary hippocampal neurons. DNA and Lipofectamine 2000 were each diluted in either BrainPhys medium for rat neurons or Neurobasal medium for mouse neurons. The DNA and Lipofectamine solutions were then combined at a 1:2 ratio and incubated for 20 minutes at room temperature. Prior to transfection, the conditioned culture medium was collected from each well and replaced with prewarmed BrainPhys or Neurobasal medium, corresponding to the species. The DNA:Lipofectamine complexes were added to the neurons, followed by a 45-minute incubation at 37°C. Subsequently, the transfection mixture was removed and the original conditioned medium was returned to the cells. All constructs were expressed for 16-24h unless otherwise stated.

### Immunocytochemistry (ICC)

The cells were fixed in 4% paraformaldehyde (PFA) + 4% (w/v) sucrose for 15 min at RT, washed with PBS and permeabilized with 0.2% Triton-X100 in PBS for 10 min. After blocking in blocking solution (2% glycine, 2% BSA, 0.2% gelatine and 50 mM NH4Cl in PBS), the primary antibody was incubated overnight at 4°C diluted in blocking solution. After three washing steps with PBS, the cells were incubated for 1 h in secondary antibody diluted in blocking solution at RT and, following another three washes with PBS, mounted in Mowiol (10% (w/v) Mowiol, 25% (v/v) glycerol, 100 mM Tris-HCl, pH 8.5, 2.5% (w/v) 1,4-Diazabicyclo[2.2.2]octane).

### Experimental methods and confocal image acquisition

#### High frequency stimulation

Electrical field stimulation was carried out using two platinum electrodes placed on the coverslip at a distance of 10mm. Cells were placed in field-stimulation chamber (RC-49MFSH; Warner Instruments, USA) and extracellular stimuli of 900 pulses (1ms; 90mA) were triggered at 10Hz frequency by an isolated pulse generator (MASTER-8, A.M.P.I., Israel) and delivered via a stimulus isolator unit (ISO-Flex, A.M.P.I.). Experiments were performed at DIV 15-16. Coverslips were either electrically stimulated or treated with tetrodotoxin (TTX, hellobio, HB1035-1, 1µM for 1h) for comparison. In the TTX-treated group, 1µM TTX was added to the neuronal growing medium (BrainPhys supplemented with SM1 and glutamine for rat neurons and Neurobasal A plus supplements for mouse neurons) for 1h and cells were kept in the incubator. Afterwards coverslips were taken out and incubated with 250µL growing medium containing TTX for 90sec. Cells corresponding to the electrically-stimulated group were stimulated in the stimulation chamber containing 250µL of neuronal growing medium for 90 sec. Cells for basal condition were not treated at all. After stimulation, all coverslips were placed in the incubator at 37 °C, 5% CO2. Incubation time was adjusted to the experimental question - e.g. for autophagy induction experiments, incubation time was 15 min, for autophagosome/ amphisome quantifications in axons, 30 min. Cells were then fixed in 4% PFA 4% saccharose for 20 minutes.

#### Live-imaging of AMPK-activity sensor

Primary neurons (DIV14-15) were transfected with AMPKAR-AV or GFP as control 48h prior before the experiment was performed. Images were acquired in a scanning confocal microscope (Leica TCS SP8 X) controlled by the software Leica LAS X, using HCX PL APO63x1.40 objective. Cells were imaged in an environmental chamber prewarmed at 37°C. The following laser lines were using for illumination: 415nm for CFP; 530nm for YFP and 561nm for Sphy-tRFP. Images were acquired in a sequential manner every 20 seconds. HFS stimulation was performed 2 minutes after baseline acquisition.

#### Application of reagents and pharmaceutical drugs

Reagents were added to the culture medium after cell silencing (in the case of TTX treatment), but prior to stimulation (for HFS condition), and remained in the medium until fixation. This procedure was applied consistently across all experiments, regardless of the experimental conditions. For Compound C, 50µM was used for 30 min incubation. SBI-0206965, was applied for 2 hours. Human TrkB Fc *bodies(500ng/m)* and TAT-peptides (30µM) were applied to the well the day before and incubated overnight. Respectively to the experiment cells were incubated 15 or 30 min. For quantum dot experiments, a coupling mix was prepared with 1µM Quantum Dots, 1µM biotin-BDNF in 1% BSA in PBS. This was vortexed for 20 minutes on low speed. 2mM of Quantum dots concentration was used in culture medium for each well. Cells were incubated 5 min prior stimulation. For stimulating the cells culture medium containing Quantum dots was used and washed away immediately after by using a quencher in tyrodes buffer. Coverslips were incubated for 30 min then fixed and processed for imaging. For experiments with dextran uptake, first TTX group was treated 1h with 1µM TTX in culture medium. Afterwards cells were incubated 5 min prior to stimulation 50µM dextran in tyrodes buffer (134 mM NaCl, 12 mM NaHCO3, 3 mM KCl, 0.34 mM Na2HPO4, 1 mM MgCl2, 10 mM HEPES, pH 7.4) with 4mM AP5 and 10mM CNOX reagents, on parafilm. For TTX group 1µM TTX was added to dextran solution and just left on parafilm. For stimulation, cells were covered in 250µl of dextran with tyrodes buffer solution during high frequency stimulation in field-stimulation chamber as described previously. Afterwards all coverslips were washed in tyrodes buffer, TTX group with TTX (1µM) respectively. Depending on the experiment, cells were incubated 10 or 30 min with 37 degrees. Independently of the treatment all cells were then fixed in 4% PFA, 4% saccharose for 20 minutes and further processed with immunocytochemistry for imaging as described previously.

Images of fixed primary neurons were acquired with an Olympus Fluoview FV3000 microscope using an oil-immersion objective (60X/1.40). Stack images were acquired with a 79x79nm pixel size, a 512x512 or 1024x1024 resolution and a Z-step of 0.30µm. Images were acquired using the 405, 488, 568 and 633 nm laser lines. Imaged axons were chosen based on specific morphological features as follows: (i) elongated, slender processes extending distally from the cell body while maintaining a uniform diameter; (ii) absence of repeated branching along the axon; and (iii) lack of any protrusions or spines along the length of the process. Image acquisition settings were optimized and kept constant for all images between groups of one data set.

#### Electron microscopy

For transmission electron microscopy (TEM) sample preparation, primary hippocampal neurons were seeded in µ-Dish 35 mm (high Grid 500, Ibidi, Germany) and kept in the incubator at 37°C and 5% CO2 until DIV14-16. Neurons were stimulated with the HFS protocol (see above) and fixed with 4% PFA (4% sucrose) in PBS overnight.

For CLEM, confocal images were acquired using Plan Apo VC 20x (NA 0.75) objective lens with a Nikon AX R CSLM.

After post-fixation with 2.5% glutaraldehyde in PBS overnight cells were washed with PBS, post-fixed with 1% OsO₄ (30 min), rinsed in PBS and ddH₂O, and stained in 1% aqueous uranyl acetate (30 min). After washing with ddH₂O, samples were ethanol-dehydrated and embedded in EPON (Carl Roth). Ultrathin sections (50nm) were cut using an Ultracut microtome (Leica Microsystems) and post-stained with 1% uranyl acetate in ethanol. Images were acquired with a FEI Tecnai G20 Twin (Thermo Fisher) at 80 kV using a 2K CCD Veleta camera (Olympus SIS).

### Metabolic labeling assays

#### Puromycilation

For puromycylation assays, neurons were incubated with 2 μM puromycin for 5 min in culture medium at 37 °C in a humidified atmosphere with 5% CO2. After Coverslips were stimulated in presence of puromycin in 250µl culture medium that was just taken out the well, respectively. Cells were incubated 10 min poststimulation and then fixed. Control cells underwent the same treatment but in the absence of puromycin. Immunocytochemistry was performed with other cell markers and prelabelled anti- puromycin-antibody.

#### FUNCAT-AHA

The FUNCAT part of the assay was performed as described previously by ref.^86^ with following modifications: Cells were preincubated 15 min in methionine-free SILAC medium supplemented with 2mM AHA then stimulated with high frequency stimulation. For experiments with Fc bodies, these were added into the culture medium the day before. Post stimulation incubation time for intensity measurements of AHA at boutons was 10 min and for FUNCAT-PLA experiments, samples were incubated 20 min. Subsequently, cells were washed two times for 5 min with PBS (1× PBS, pH 7.4) and fixed for 15 min in PFA-sucrose (4% paraformaldehyde, 4% sucrose in PBS) at room temperature, washed, permeabilized with 0.5% Triton X-100 in 1× PBS, pH 7.4, for 10 min and blocked with blocking buffer for 1 h. Afterwards cells were equilibrated by two washes with 1 x PBS (pH 7.8) to start CuAAC click reaction.

The reducing agent tris(2-carboxyethyl)phosphine (TCEP) in combination with CuSO4 was used to generate the Cu(I) catalyst for CuAAC to avoid copper bromide-derived precipitates. A click reaction mix composed of 200 μM triazole ligand Tris[(1-benzyl-1H-1,2,3-triazol-4-yl)methyl]amine (TBTA), 25 μM biotin alkyne tag, 500 μM TCEP and 200 μM CuSO4 was mixed in PBS, pH 7.8. Every reagent was added step by step with intense vortexing in between. The click reaction mixture was prepared immediately before application to the cells and CuAAC was performed overnight at room temperature and then washed with PBS and 0.5 % Triton in PBS, permeabilized and further processed eather as described in the sections ‘FUNCAT-PLA’ or ‘immunocytochemistry’.

#### FUNCAT-Proximity ligation assay (PLA)

For proximity ligation assay, anti-biotin in combination with protein-specific antibodies were used. Detection was carried out with Duolink reagents according to manufacturer’s protocol with slight modifications to dilutions. For secondary antibodies, Duolink rb PLAplus and ms PLAminus probes were used. The ‘Duolink Detection reagents Red’ were used for ligation, amplification and label probe binding.

Briefly, after metabolic labeling, permeabilization and washing (see section ‘FUNCAT’) cells were blocked in blocking buffer (4 % goat serum in PBS, 1 h) and incubated with primary antibody pairs diluted in blocking buffer (1.5 h at room temperature). After washing, PLA probes were applied in 1:10 dilution in blocking buffer for 1 h at 37 °C, washed several times with wash buffer A (0.01 M Tris, 0.15 M NaCl, 0.05 % Tween20) and incubated for 30 min with the ligation reaction containing the circularization oligos and T4 ligase prepared according to the manufacturer’s recommendations (Duolink Detection reagents Red, Sigma) in a prewarmed humidified chamber at 37 °C. Amplification and label probe binding was performed after further washes with wash buffer A with the amplification reaction mixture containing Phi29 Polymerase and the fluorophore-labeled detection oligo prepared according to the manufacturer’s recommendations in a prewarmed humidified chamber at 37 °C for 100 min. Amplification was stopped by three washes in 0.2 M Tris, 0.1 M NaCl pH 7.5 followed by washes in PBS pH 7.4. For better signal stability, cells were postfixed for 10 min at room temperature in PFA-sucrose, washed with PBS and processed further for immunohistochemistry. After staining, imaging, quantification and analysis was conducted as described in previous section.

### Nanoscopy

#### STED

Stimulated emission depletion (STED) imaging was carried out using an Abberior multichannel confocal/STED microscope based on a Nikon Ti-E microscope body with perfect focus system controlled by Imspector Software (Abberior Instruments). An x60 P-Apo oil immersion objective (NA 1.4) was used. STED laser lines 561 and 640nm were used for excitation and a laser line at 775nm was used for fluorescence depletion.

#### DNA-PAINT/MINFLUX

Primary neurons at DIV14-16 were stimulated using HFS and kept in the incubator at 37°C 5%CO2 for 30 minutes. After fixation with 4% PFA, immunocytochemistry was performed as described above using mouse-anti LC3 (1:300; MBL) and rabbit anti-pTrk (1:300). Samples were incubated with the following secondary antibodies for one hour at room temperature: anti-mouse 488 (1:500); anti-mouse MF4 (1:200); anti-rabbit MF5 (1:500). Following secondary-antibody labeling, coverslips were washed with PBS for 3 times and kept in PBS at 4°C until the experiment was performed.

Sample preparation before imaging was performed using the MASSIVE-DNA-PAINT kits optimized for MINFLUX microscopy (Abberior) according to manufactureŕs instruction. Gold nanorods (Nanopartz Inc) were used as fiducials and were applied to the coverslips for 5 minutes followed by a 15-minute treatment with poly-L-lysine. Right before imaging, samples were incubated with imager strands (MF4 1:1000; MF5 1/2000) which were diluted in PBS, and mounted on dimple slides sealed with two-part silicon glue for easy imaging and storage. MINFLUX nanoscopy, corresponding confocal imaging as well as image rendering were performed in a MINFLUX setup as previously described^36^.

Following acquisition, single-molecule localizations were filtered. Localizations that were found in >4 scans were taken into consideration. Images were visualized using Fiji and IMARIS.

### In-situ hybridization and ExFISH

#### RNA-ISH in primary neurons

ISH was performed in PFA-fixed primary murine neurons using RNAscope™ Multiplex Fluorescent Reagent Kit v2 (ACD, Bio-techne) as instructed by the manufacturer’s user manual. In brief, cells were fixed in 4% PFA + 0.1M sucrose for 15min at RT and washed three times with PBS. The cells were permeabilized at RT in 0.2 % Triton-X100, followed by a 10min incubation in H2H2 (pretreatment reagent) and two washes in Millipore water. As final pretreatment, the cells were incubated in RNAscope-protease-III (1:30 dilution) for 10min at RT, followed by two PBS washes. The RNAscope probes were hybridized for 2 h at 40°C in the hybEZ oven system (ACD). Bassoon transcripts were detected with a custom-made RNAscope probe targeting the region encoding amino acids 2468–3128, while for the remaining transcripts commercially available probes were used. Amplification steps were performed as described in the manual, followed by the dye incubation using the TSA Vivid™ Fluorophore Kit 570 (Tocris, #7526) with a dilution of 1:2000 for RNAscope and 1:1000 for ExFISH. The ISH was completed using the RNAscope Multiplex FL v2 HRP blocker solution for 15 min at 40°C and two final washes in RNAscope wash buffer. RNAscope-labelled coverslips were used for Co-Immunocytochemistry or ExFISH experiments.

#### ExFISH

RNAscope-labelled and already co-immunostained coverslips were used for ExFISH experiments. Each Bsn, Syn, and Mapt ISH-labelled coverslip was protein-costained for Synapsin1/2 (SySy, 106 004) to label presynaptic boutons and MAP2 (EnCor, CPCA-MAP2) to orient in the gel. Abberior STAR green and STAR red were used as secondary antibodies at a dilution of 1:100. ExFISH was performed as previously described (citations) with the following modifications: RNAscope-labelled coverslips were anchored as described in ref.^87^, starting with RNA-anchoring (LabelX O/N at RT) and followed by protein-anchoring (AcX O/N at RT). Gel polymerization was performed as described in (Delling), followed by homogenization in digestion buffer (1 M NaCl, 1 mM EDTA, 0.5 % Triton X-100, 50 mM Tris with 8 U/mL of proteinase K for 5 hrs at 37°C, slowly shaking. Gels were detached from the coverslips and washed three times in PBS at RT. Gels were measured, cut, and transferred in a 12-well plate and the protein co-immunostaining was repeated with a primary antibody incubation time from 48-72 hrs at 4°C. After secondary antibody incubation and final PBS washes, gels were transferred to 10cm petri dishes, expanded and mounted.

#### RNAscope image acquisition and quantification

RNAscope images were acquired using a Leica TCS SPE II confocal microscope (Wetzlar, Germany) and analyzed using FIJI (ImageJ 2.16). For colocalization analysis, a bouton mask was generated using the thresholding function and ISH signals within the mask were counted. Unpaired Student’s t test was used for statistical comparision.

#### Expansion image acquisition

ExFISH images were acquired on a STEDYCON module (Abberior) fixed to an Axio Imager.Z2 (Zeiss) equipped using a 100x/1.30 (Zeiss) oil-immersion objective in confocal mode. Microscope settings were set using secondary antibody controls for protein channels and RNAscope negative control probes (3-plex Negative Ctrl, Cat No. 320871, ACD) for ISH channels as baseline.

### Image and statistical analysis

The analysis of images was conducted using the free software Fiji (ImageJ). Maximum intensity projections of the raw image stacks were generated for quantification. For intensity measurements, the thresholding function in ImageJ was applied to define regions corresponding to presynaptic antibody labeling. Integrated density values within these regions were quantified to compare TTX and stimulated groups. For colocalization analysis, the thresholding function was first used to mask imaged axons, followed by identification of signals from labeled proteins of interest. Consistent threshold values were applied across the same channels in different experimental groups. Regions of interest (ROIs) were counted using the multi-point tool. Colocalization events were defined by the overlap of ROIs from different channels and saved as RoiSets.

Statistical Analysis was performed with GraphPad Prism 10. Unless otherwise stated, simple t-test was used to compare 2 unpaired samples and One-way ANOVA was used to compare more than two groups. For analysis of wildtype/ko animals, as well as AMPK-DN assays, two-way ANOVA was conducted. Use of asterisks: p-value: <0.0001 to 0.001 ***, 0.001 to 0.01 **, 0.01 to 0.05 *, ≥ 0.05 not significant (ns).

Data in the manuscript are shown as mean ± SEM and *n* numbers used for statistics are depicted in each panel or figure legends. Graphs and statistical analysis were made with GraphPad Prism (GraphPad Software).

---

**Extended data figure 1.**
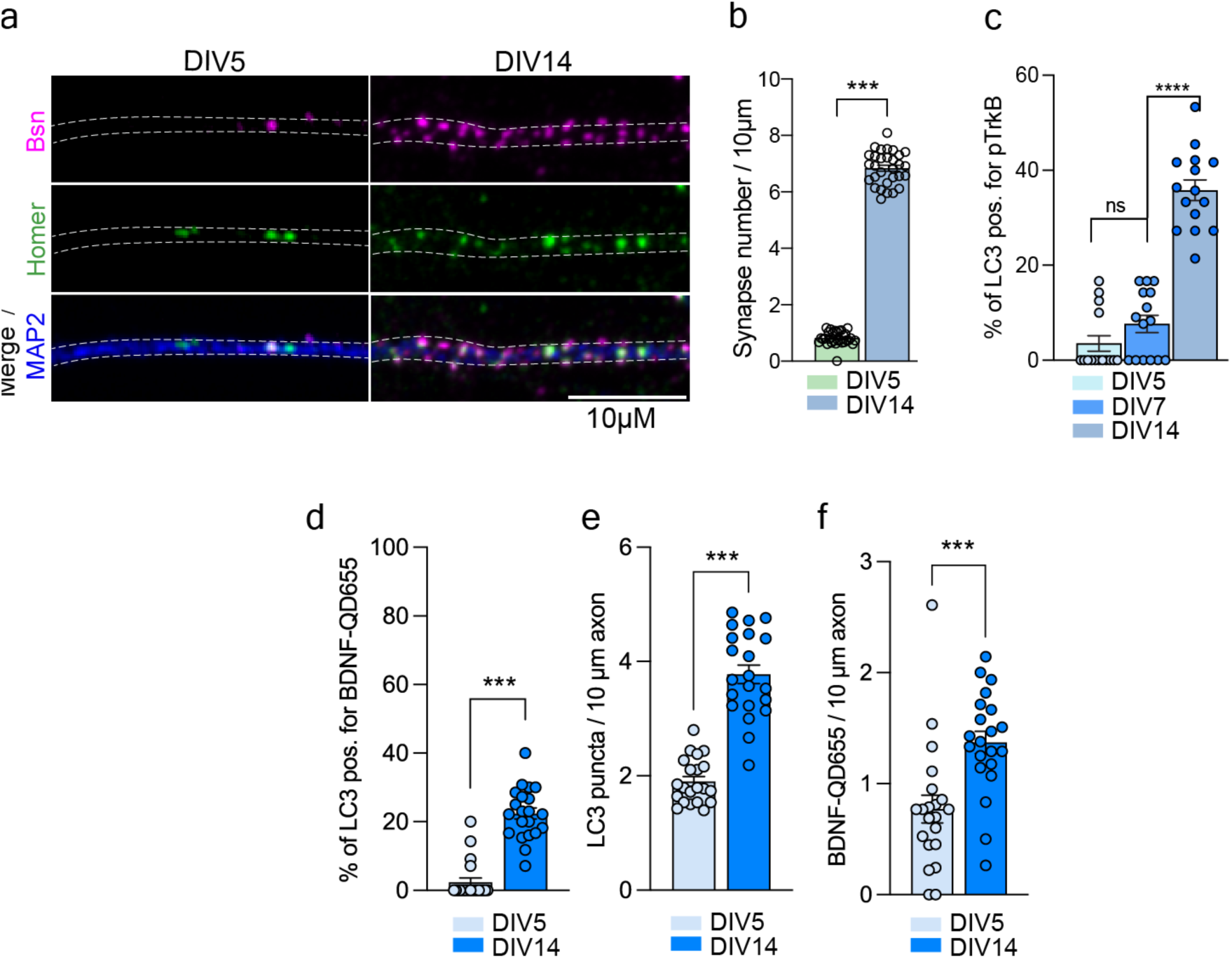
Amphisome biogenesis occurs in mature neurons. A) Representative images of neurons at DIV5 and 14 immunostained against MAP2 as neurite marker, and Bassoon and Homer as synaptic markers. Scale bar indicates 10µm. B) Quantification of synapse number from S1A). (*n=30 from 3 independent cultures)* Belongs to Fig.1F; C) Percentage of amphisomes from total LC3-vesicles analyzed in axons from cells at DIV5, DIV7 and DIV14. (n = 15 from 2 independent cultures) D) Percentage of all quantified LC3 puncta colocalizing with BDNF in axons under basal conditions in DIV5 and DIV14 hippocampal neurons. Fig 1H+I E) Quantification of LC3 puncta in 10 µm axon from the experiment in C/Fig 1H+I F) Quantification of BDNF-QD655 in 10 µm axon from the experiment in C/Fig 1H+I. (D-F) *n=21 from 3 independent cultures* Data are plotted as average ± SEM. ^∗∗∗^ represents p < 0.0001 to 0.001, ^∗^ represents p < 0.01 to 0.05 by two-tailed Mann-Whitney U test.

**Extended data figure 2.**
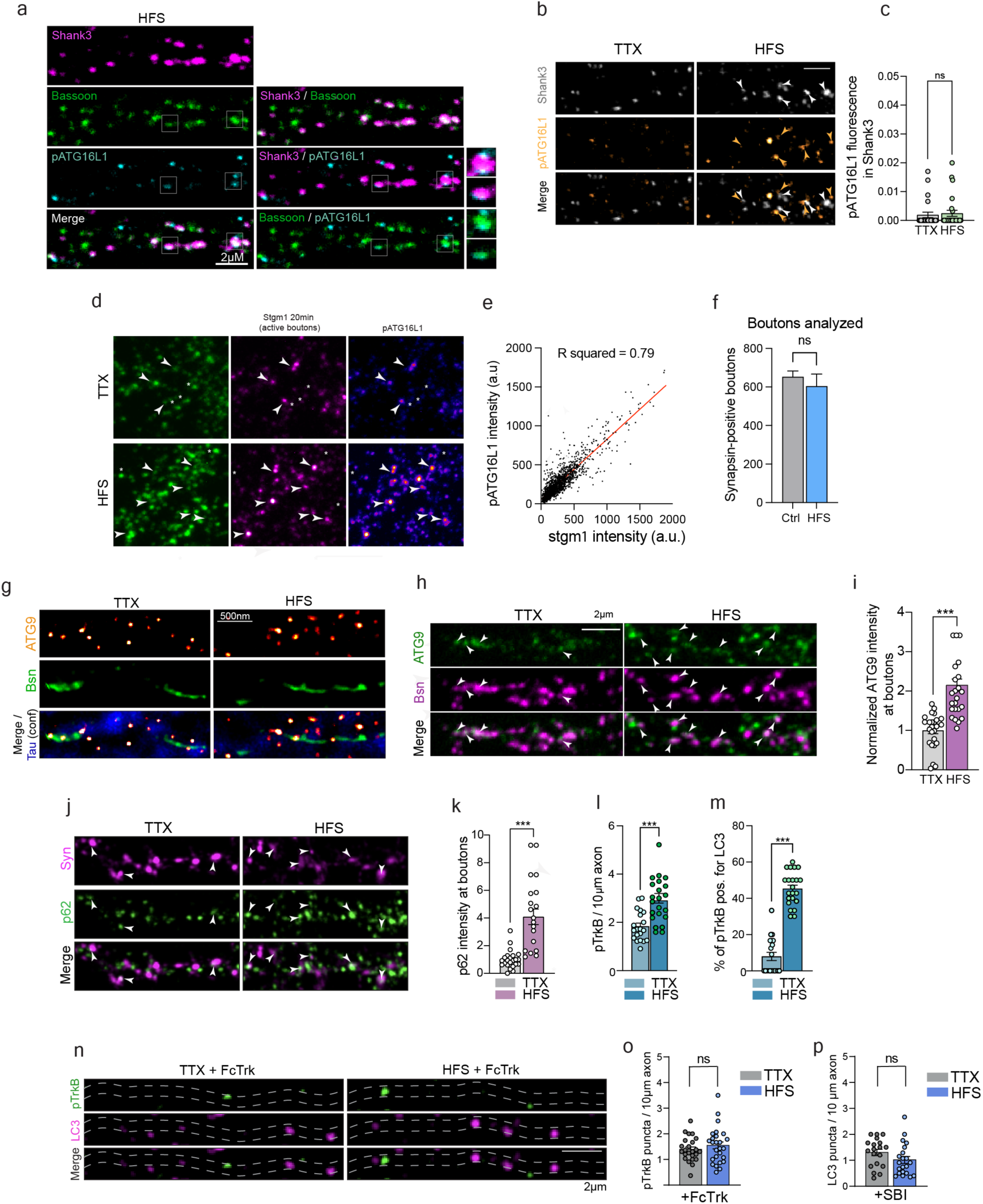
Electrical field stimulation induces autophagy activation at boutons. A) Representative images of TTX or HFS-treated primary neurons immunostained for Shank3, Bassoon and pATG16L1. B) Representative images of TTX or HFS-treated cells immunostained against pATG16L1 and Shank3. Scale bar 2µm. C) Quantification of pATG16L1 intensity measured at postsynaptic sites labelled with Shank3 in TTX and HFS neurons. (n=24 (TTX); n=27(HFS) from 3 independent cultures) D) Representative image of control primary neurons or HFS in the presence of an anti-synaptotagmin1 (luminal domain) antibody whose epitope is only exposed upon fusion of SV with the plasma membrane. After fixation cells were immunostained against pATG16L1. E) Correlation analysis of pATG16l1 intensity to Stgm1 intensity in synapsin mask. *Pearson correlation analysis was performed and best-fit linear regression line is plotted. R squared = 0.79*. F) Bar graph shows the total number of analyzed boutons for the experiment in C in the HFS and control group. G) 2D STED image of ATG9 and Bassoon in axons, labelled in blue (confocal) with Tau. Scale bar is 500nm. H) Representative images of the presynapse labelled with Bassoon and ATG9. I) Quantification of ATG9 intensity measured in Bassoon-positive boutons from TTX and HFS groups. (n=27 from 3 independent cultures) J) Representative images showing immunostaining of primary neurons against p62 and Synapsin as presynaptic marker for TTX and HFS conditions. K) Bar graph depicts p62 intensity measured at presynapses detected by Synapsin comparing TTX and HFS conditions. *n=21 (FS) n=23 (TTX) from 3 independent cultures* L) Quantification of pTrkB puncta in axons. TTX and HFS compared. To Fig2,P. M) Percentage of pTrkB puncta positive for LC3 in axons. To Fig2,P). (L+M) *n=22 (FS) n=20 (TTX) from 3 independent cultures*. N) Representative images of primary neurons incubated with Fc-Trk to chelate endogenous BDNF and immunostained against amphisome markers following TTX or HFS treatment. O) Quantification of pTrk puncta pero 10µm axons from the experiment in N. P) Quantification of LC3 puncta from the experiment in Fig 2P in the presence of SBI-0206965 for 2h. Data are plotted as average ± SEM. ^∗∗∗^ represents p < 0.0001 to 0.001 by two-tailed Mann-Whitney U test.

**Extended data figure 3.**
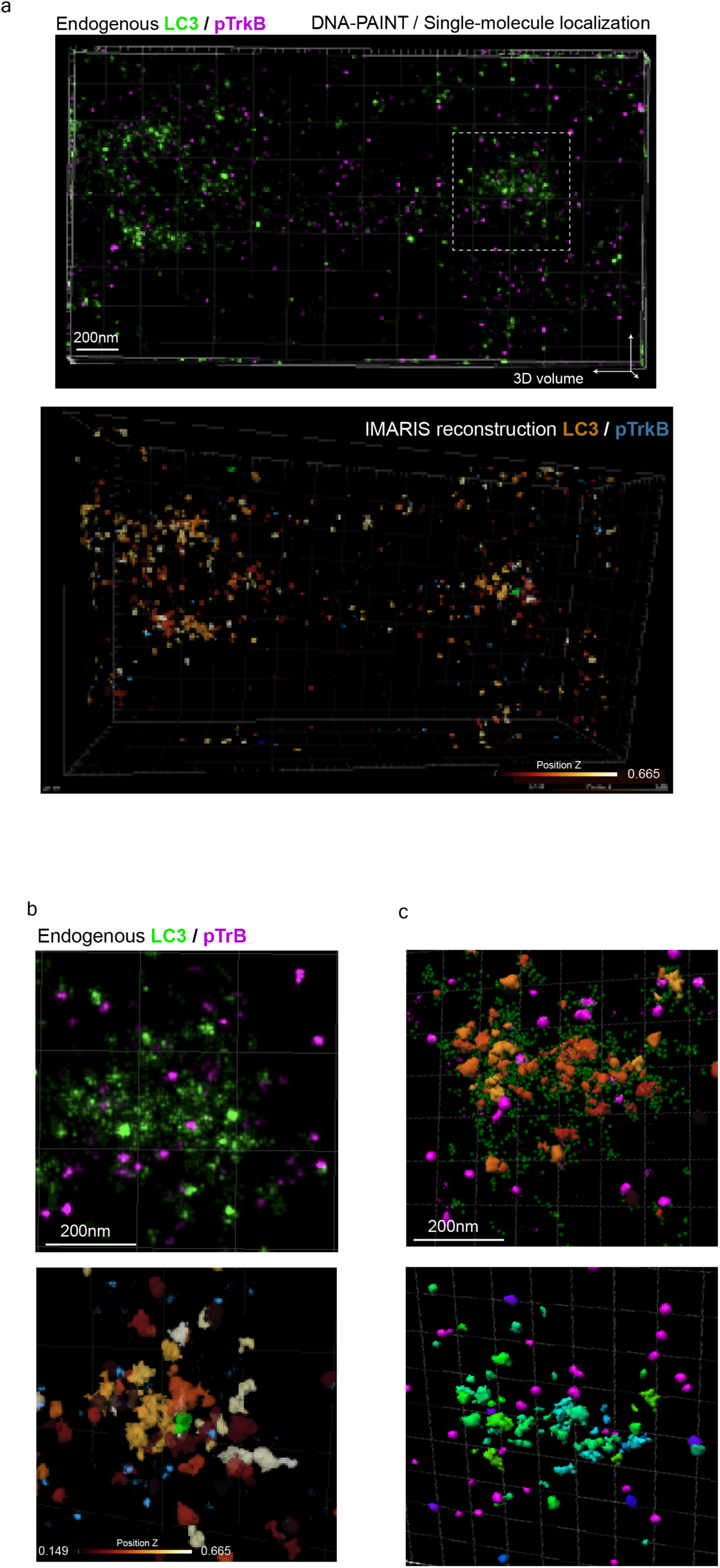
Single molecule localization of endogenous stained LC3 and pTrkB. A) 3D volume of single molecule localizations obtained for endogenous LC3 and pTrkB with DNA-PAINT/MINFLUX. Scale bar is 200nm B) Zoomed image of LC3 and pTrkB 3D localization from ROI (scattered square) shown in A. The image below show a colour-coded reconstruction. Scale bar is 200nm. C) A different example from another experiment of an amphisome images by DNA-PAINT/MINFLUX.

**Extended data figure 4.**
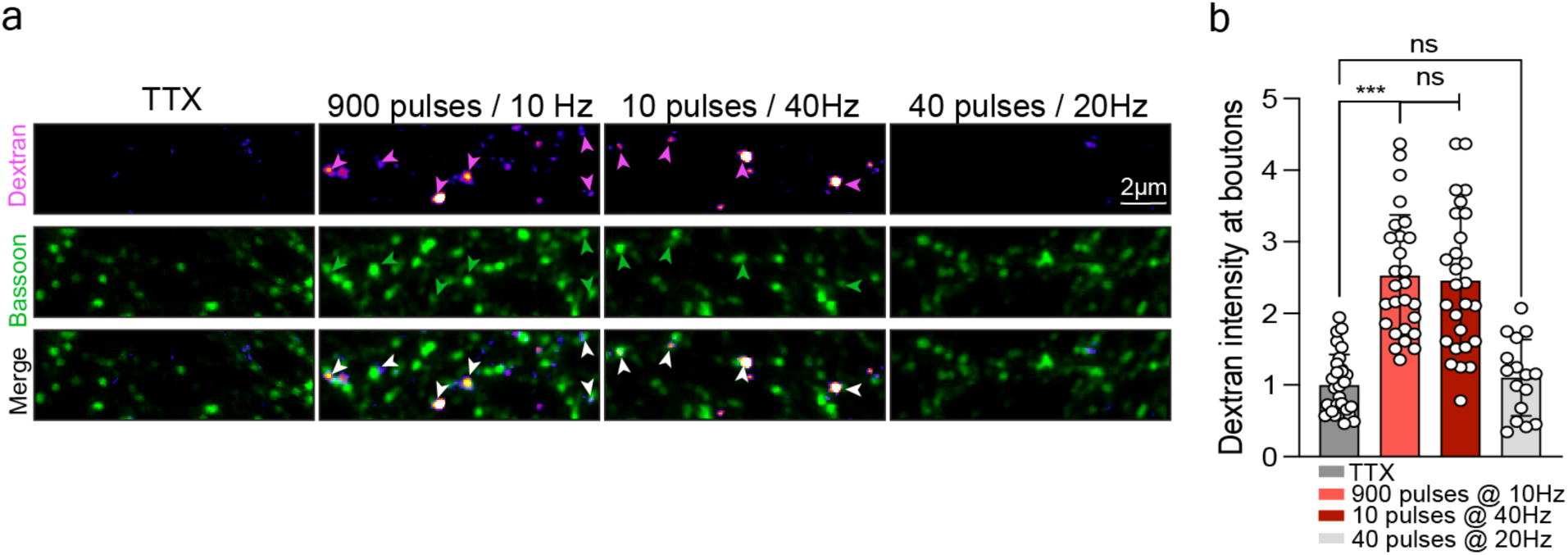
Amphisomes are formed from bulk-endosomes. A) Representative images showing dextran at puncta detected by Bassoon staining following different electrical stimulation protocols. B) Bar graph shows dextran intensity at boutons from the experiment in B. TTX group compared to different stimulation protocols. 900 pulses@10Hz used in this study was compared to 10 pulses@40Hz employed in ref.^39^ to induce ABDE (positive control) and to 40 pulses@20Hz employed by others^90^ to mobilize the ready-releasable pool of SV (negative control). n= 29 (TTX); n=27(HFS); 27 (10 pulses, 40Hz); n=16 (40 pulses, 20Hz) Data are presented as average ± SEM. Statistical analysis was performed using the Kruskal–Wallis test followed by Dunn’s multiple-comparisons test. ^∗∗∗^ represents p < 0.0001 to 0.001.

**Extended data figure 5.**
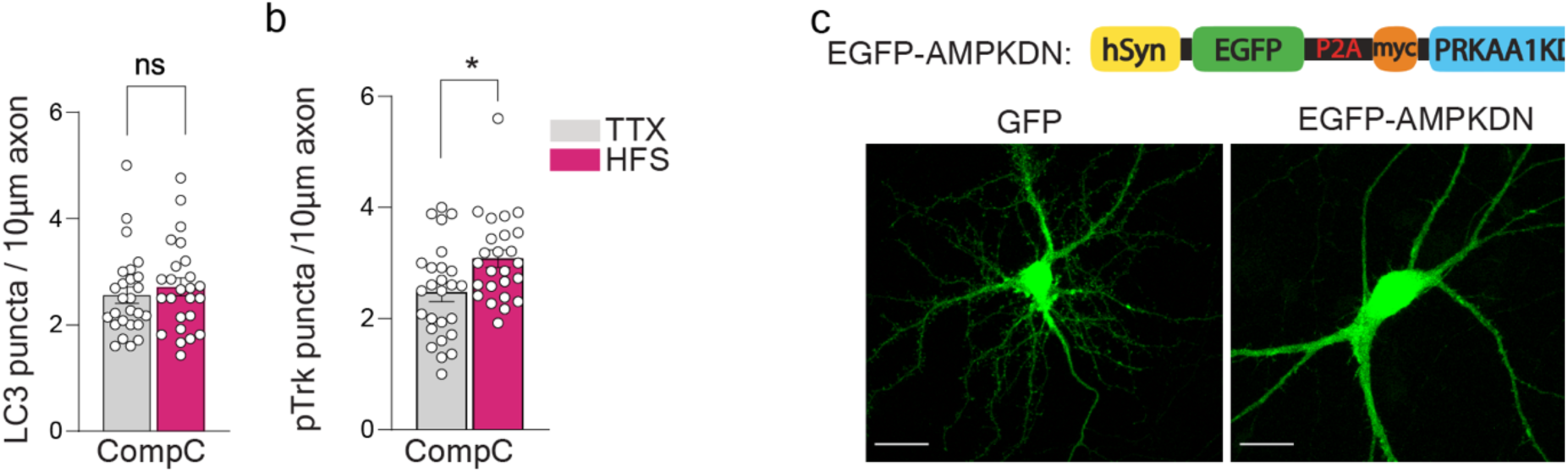
AMPK inhibition prevents synaptic autophagy. A) Quantification of LC3 puncta per 10µm axon from the experiment in Fig 5G. B) Quantification of pTrkB puncta per 10µm from the experiment in G. C) Scheme showing the expression vector used in AMPK-dominant negative (AMPK-DN) experiments. Expression of GFP and AMPK-DN in the same cells is achieved by means of a P2A sequence. In control experiments an empty GFP construct is used. Representative images of transfected cells are shown below. Scale bar indicates 20µm. *(A+B) n=25(HFS+Comp C.); n=26 (TTX+Comp. C) from 3 independent cultures; Data are plotted as average ± SEM. ^∗^ represents p = 0.01 to 0.05.by two-tailed Mann-Whitney U test*.

**Extended Data Figure 6.**
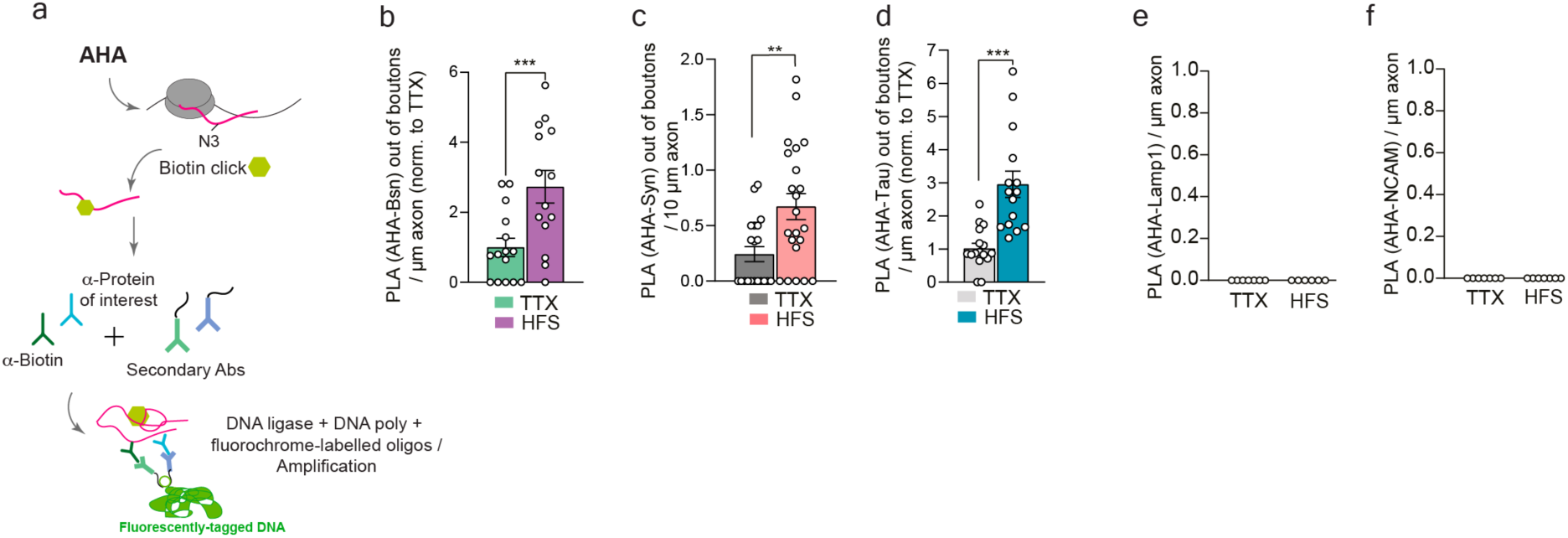
Local protein synthesis in axons. A) Schematic representation depicting the principle of FUNCAT-PLA used in the experiments from B-F and Fig8 G-W. B) Quantification of PLA signal for Bsn along axons but outside of boutons labelled with Piccolo per 10µm of axon from the experiment in Fig8 G-I. n=22(TTX); n=19(HFS) from 3 independent cultures C) Quantification of PLA signal for Syn along axons but outside of boutons labelled with Piccolo per 10µm of axon from the experiment in Fig8 J-L. n=20(TTX); n=22(HFS) from 3 independent cultures D) Quantification of PLA signal for Tau along axons but outside of boutons labelled with Piccolo per 10µm of axon from the experiment in Fig8 M-O. n=22(TTX); n=19(HFS) from 3 independent cultures E-F) Quantification of PLA signal of locally synthesized LAMP1 (E; n=19 from 3 independent cultures) or NCAM (F; n=19(TTX); n=20(HFS) from 3 independent cultures). Data are presented as average ± SEM. Statistical analysis was performed by two-tailed Mann-Whitney U test. ^∗∗∗^ represents p < 0.0001 to 0.001, ** shows p= 0.001 to 0.01.

## References

1. Biever, A. et al. Monosomes actively translate synaptic mRNAs in neuronal processes. Science 367 (2020).

2. Hafner, A.S., Donlin-Asp, P.G., Leitch, B., Herzog, E. & Schuman, E.M. Local protein synthesis is a ubiquitous feature of neuronal pre- and postsynaptic compartments. Science 364 (2019).

3. Rankovic, B. et al. RNA promotes synapsin coacervation and modulates local translation. bioRxiv (2025).

4. Ackermann, F., Waites, C.L. & Garner, C.C. Presynaptic active zones in invertebrates and vertebrates. EMBO Rep 16, 923–938 (2015).

5. Schoch, S. & Gundelfinger, E.D. Molecular organization of the presynaptic active zone. Cell Tissue Res 326, 379–391 (2006).

6. Siksou, L. et al. Three-dimensional architecture of presynaptic terminal cytomatrix. J Neurosci 27, 6868–6877 (2007).

7. Sudhof, T.C. The presynaptic active zone. Neuron 75, 11–25 (2012).

8. Wilhelm, B.G. et al. Composition of isolated synaptic boutons reveals the amounts of vesicle trafficking proteins. Science 344, 1023–1028 (2014).

9. Milovanovic, D. & De Camilli, P. Synaptic Vesicle Clusters at Synapses: A Distinct Liquid Phase? Neuron 93, 995–1002 (2017).

10. Milovanovic, D., Wu, Y., Bian, X. & De Camilli, P. A liquid phase of synapsin and lipid vesicles. Science 361, 604–607 (2018).

11. Karpova, A. et al. Neuronal autophagy in the control of synapse function. Neuron 113, 974–990 (2025).

12. Andres-Alonso, M., Kreutz, M.R. & Karpova, A. Autophagy and the endolysosomal system in presynaptic function. Cell Mol Life Sci 78, 2621–2639 (2021).

13. Grochowska, K.M., Andres-Alonso, M., Karpova, A. & Kreutz, M.R. The needs of a synapse-How local organelles serve synaptic proteostasis. EMBO J 41, e110057 (2022).

14. Stavoe, A.K.H. & Holzbaur, E.L.F. Autophagy in Neurons. Annu Rev Cell Dev Biol 35, 477–500 (2019).

15. Kuijpers, M. et al. Neuronal Autophagy Regulates Presynaptic Neurotransmission by Controlling the Axonal Endoplasmic Reticulum. Neuron 109, 299–313 e299 (2021).

16. Wosnitzka, E., Gambarotto, L. & Nikoletopoulou, V. Macroautophagy at the service of synapses. Curr Opin Neurobiol 93, 103054 (2025).

17. Egan, D.F. et al. Phosphorylation of ULK1 (hATG1) by AMP-activated protein kinase connects energy sensing to mitophagy. Science 331, 456–461 (2011).

18. Feng, Y. et al. Interplay of energy metabolism and autophagy. Autophagy 20, 4–14 (2024).

19. Mao, K. & Klionsky, D.J. AMPK activates autophagy by phosphorylating ULK1. Circ Res 108, 787–788 (2011).

20. Andres-Alonso, M. et al. SIPA1L2 controls trafficking and local signaling of TrkB-containing amphisomes at presynaptic terminals. Nat Commun 10, 5448 (2019).

21. Kononenko, N.L. et al. Retrograde transport of TrkB-containing autophagosomes via the adaptor AP-2 mediates neuronal complexity and prevents neurodegeneration. Nat Commun 8, 14819 (2017).

22. Kulkarni, V.V., Stempel, M.H., Anand, A., Sidibe, D.K. & Maday, S. Retrograde Axonal Autophagy and Endocytic Pathways Are Parallel and Separate in Neurons. J Neurosci 42, 8524–8541 (2022).

23. Sidibe, D.K. et al. Brain-derived neurotrophic factor stimulates the retrograde pathway for axonal autophagy. J Biol Chem 298, 102673 (2022).

24. Fernandez-Alfonso, T. & Ryan, T.A. The kinetics of synaptic vesicle pool depletion at CNS synaptic terminals. Neuron 41, 943–953 (2004).

25. Kwon, S.E. & Chapman, E.R. Synaptophysin regulates the kinetics of synaptic vesicle endocytosis in central neurons. Neuron 70, 847–854 (2011).

26. Virmani, T., Atasoy, D. & Kavalali, E.T. Synaptic vesicle recycling adapts to chronic changes in activity. J Neurosci 26, 2197–2206 (2006).

27. Lie, P.P.Y. et al. Post-Golgi carriers, not lysosomes, confer lysosomal properties to pre-degradative organelles in normal and dystrophic axons. Cell Rep 35, 109034 (2021).

28. Soukup, S.F. et al. A LRRK2-Dependent EndophilinA Phosphoswitch Is Critical for Macroautophagy at Presynaptic Terminals. Neuron 92, 829–844 (2016).

29. Gundelfinger, E.D., Karpova, A., Pielot, R., Garner, C.C. & Kreutz, M.R. Organization of Presynaptic Autophagy-Related Processes. Front Synaptic Neurosci 14, 829354 (2022).

30. Tian, W. et al. An antibody for analysis of autophagy induction. Nat Methods 17, 232–239 (2020).

31. Alsaadi, R.M. et al. ULK1-mediated phosphorylation of ATG16L1 promotes xenophagy, but destabilizes the ATG16L1 Crohn’s mutant. EMBO Rep 20, e46885 (2019).

32. Dikic, I. & Elazar, Z. Mechanism and medical implications of mammalian autophagy. Nat Rev Mol Cell Biol 19, 349–364 (2018).

33. Hamaoui, D. & Subtil, A. ATG16L1 functions in cell homeostasis beyond autophagy. FEBS J 289, 1779–1800 (2022).

34. Kraszewski, K. et al. Synaptic vesicle dynamics in living cultured hippocampal neurons visualized with CY3-conjugated antibodies directed against the lumenal domain of synaptotagmin. J Neurosci 15, 4328–4342 (1995).

35. Magne, J. & Green, D.R. LC3-associated endocytosis and the functions of Rubicon and ATG16L1. Sci Adv 8, eabo5600 (2022).

36. Schmidt, R. et al. MINFLUX nanometer-scale 3D imaging and microsecond-range tracking on a common fluorescence microscope. Nat Commun 12, 1478 (2021).

37. Clayton, E.L. & Cousin, M.A. The molecular physiology of activity-dependent bulk endocytosis of synaptic vesicles. J Neurochem 111, 901–914 (2009).

38. Clayton, E.L., Evans, G.J. & Cousin, M.A. Bulk synaptic vesicle endocytosis is rapidly triggered during strong stimulation. J Neurosci 28, 6627–6632 (2008).

39. Nicholson-Fish, J.C., Kokotos, A.C., Gillingwater, T.H., Smillie, K.J. & Cousin, M.A. VAMP4 Is an Essential Cargo Molecule for Activity-Dependent Bulk Endocytosis. Neuron 88, 973–984 (2015).

40. Anggono, V. et al. Syndapin I is the phosphorylation-regulated dynamin I partner in synaptic vesicle endocytosis. Nat Neurosci 9, 752–760 (2006).

41. Clayton, E.L. et al. The phospho-dependent dynamin-syndapin interaction triggers activity-dependent bulk endocytosis of synaptic vesicles. J Neurosci 29, 7706–7717 (2009).

42. Li, S. & Sheng, Z.H. Energy matters: presynaptic metabolism and the maintenance of synaptic transmission. Nat Rev Neurosci 23, 4–22 (2022).

43. Li, S., Xiong, G.J., Huang, N. & Sheng, Z.H. The cross-talk of energy sensing and mitochondrial anchoring sustains synaptic efficacy by maintaining presynaptic metabolism. Nat Metab 2, 1077–1095 (2020).

44. Monday, H.R., Kharod, S.C., Yoon, Y.J., Singer, R.H. & Castillo, P.E. Presynaptic FMRP and local protein synthesis support structural and functional plasticity of glutamatergic axon terminals. Neuron 110, 2588–2606 e2586 (2022).

45. Ostroff, L.E. et al. Axon TRAP reveals learning-associated alterations in cortical axonal mRNAs in the lateral amgydala. Elife 8 (2019).

46. Shigeoka, T. et al. Dynamic Axonal Translation in Developing and Mature Visual Circuits. Cell 166, 181–192 (2016).

47. Younts, T.J. et al. Presynaptic Protein Synthesis Is Required for Long-Term Plasticity of GABA Release. Neuron 92, 479–492 (2016).

48. Castillo, P.E., Jung, H., Klann, E. & Riccio, A. Presynaptic Protein Synthesis in Brain Function and Disease. J Neurosci 43, 7483–7488 (2023).

49. Fonkeu, Y. et al. How mRNA Localization and Protein Synthesis Sites Influence Dendritic Protein Distribution and Dynamics. Neuron 103, 1109–1122 e1107 (2019).

50. Rangaraju, V., Lauterbach, M. & Schuman, E.M. Spatially Stable Mitochondrial Compartments Fuel Local Translation during Plasticity. Cell 176, 73–84 e15 (2019).

51. Rangaraju, V., tom Dieck, S. & Schuman, E.M. Local translation in neuronal compartments: how local is local? EMBO Rep 18, 693–711 (2017).

52. Costa, R.O., Martins, L.F., Tahiri, E. & Duarte, C.B. Brain-derived neurotrophic factor-induced regulation of RNA metabolism in neuronal development and synaptic plasticity. Wiley Interdiscip Rev RNA 13, e1713 (2022).

53. Sahoo, P.K., Smith, D.S., Perrone-Bizzozero, N. & Twiss, J.L. Axonal mRNA transport and translation at a glance. J Cell Sci 131 (2018).

54. Cioni, J.M. et al. Late Endosomes Act as mRNA Translation Platforms and Sustain Mitochondria in Axons. Cell 176, 56–72 e15 (2019).

55. Okerlund, N.D. et al. Bassoon Controls Presynaptic Autophagy through Atg5. Neuron 93, 897–913 e897 (2017).

56. Reshetniak, S. et al. The synaptic vesicle cluster as a controller of pre- and postsynaptic structure and function. J Physiol (2024).

57. Hoffmann, C. et al. Synapsin condensation controls synaptic vesicle sequestering and dynamics. Nat Commun 14, 6730 (2023).

58. Longfield, S.F. et al. Tau forms synaptic nano-biomolecular condensates controlling the dynamic clustering of recycling synaptic vesicles. Nat Commun 14, 7277 (2023).

59. Qiu, H. et al. Short-distance vesicle transport via phase separation. Cell 187, 2175–2193 e2121 (2024).

60. Glock, C. et al. The translatome of neuronal cell bodies, dendrites, and axons. Proc Natl Acad Sci U S A 118 (2021).

61. Yperman, K. & Kuijpers, M. Neuronal endoplasmic reticulum architecture and roles in axonal physiology. Mol Cell Neurosci 125, 103822 (2023).

62. Hernandez-Diaz, S., et al. Synaptogyrin regulates neuronal activity dependent autophagy to degrade synaptic vesicle components and pathological Tau. bioRxiv (2023).

63. Azarnia Tehran, D., Kuijpers, M. & Haucke, V. Presynaptic endocytic factors in autophagy and neurodegeneration. Curr Opin Neurobiol 48, 153–159 (2018).

64. Soukup, S.F. & Verstreken, P. EndoA/Endophilin-A creates docking stations for autophagic proteins at synapses. Autophagy 13, 971–972 (2017).

65. Vijayan, V. & Verstreken, P. Autophagy in the presynaptic compartment in health and disease. J Cell Biol 216, 1895–1906 (2017).

66. Boucrot, E. et al. Endophilin marks and controls a clathrin-independent endocytic pathway. Nature 517, 460–465 (2015).

67. Milosevic, I. et al. Recruitment of endophilin to clathrin-coated pit necks is required for efficient vesicle uncoating after fission. Neuron 72, 587–601 (2011).

68. Watanabe, S. et al. Synaptojanin and Endophilin Mediate Neck Formation during Ultrafast Endocytosis. Neuron 98, 1184–1197 e1186 (2018).

69. Murdoch, J.D. et al. Endophilin-A Deficiency Induces the Foxo3a-Fbxo32 Network in the Brain and Causes Dysregulation of Autophagy and the Ubiquitin-Proteasome System. Cell Rep 17, 1071–1086 (2016).

70. Vanhauwaert, R. et al. The SAC1 domain in synaptojanin is required for autophagosome maturation at presynaptic terminals. EMBO J 36, 1392–1411 (2017).

71. Kononenko, N.L. et al. Clathrin/AP-2 mediate synaptic vesicle reformation from endosome-like vacuoles but are not essential for membrane retrieval at central synapses. Neuron 82, 981–988 (2014).

72. McMahon, H.T. & Boucrot, E. Molecular mechanism and physiological functions of clathrin-mediated endocytosis. Nat Rev Mol Cell Biol 12, 517–533 (2011).

73. Imai, K. et al. Atg9A trafficking through the recycling endosomes is required for autophagosome formation. J Cell Sci 129, 3781–3791 (2016).

74. Sawa-Makarska, J. et al. Reconstitution of autophagosome nucleation defines Atg9 vesicles as seeds for membrane formation. Science 369 (2020).

75. Yang, S. et al. Presynaptic autophagy is coupled to the synaptic vesicle cycle via ATG-9. Neuron (2022).

76. Cagnetta, R., Flanagan, J.G. & Sonenberg, N. Control of Selective mRNA Translation in Neuronal Subcellular Compartments in Health and Disease. J Neurosci 43, 7247–7263 (2023).

77. Bourke, A.M., Schwarz, A. & Schuman, E.M. De-centralizing the Central Dogma: mRNA translation in space and time. Mol Cell 83, 452–468 (2023).

78. Garza-Lombo, C., Schroder, A., Reyes-Reyes, E.M. & Franco, R. mTOR/AMPK signaling in the brain: Cell metabolism, proteostasis and survival. Curr Opin Toxicol 8, 102–110 (2018).

79. Roach, P.J. AMPK -> ULK1 -> autophagy. Mol Cell Biol 31, 3082–3084 (2011).

80. Dieni, S. et al. BDNF and its pro-peptide are stored in presynaptic dense core vesicles in brain neurons. J Cell Biol 196, 775–788 (2012).

81. Heyden, A. et al. Hippocampal enlargement in Bassoon-mutant mice is associated with enhanced neurogenesis, reduced apoptosis, and abnormal BDNF levels. Cell Tissue Res 346, 11–26 (2011).

82. Hoffmann-Conaway, S. et al. Parkin contributes to synaptic vesicle autophagy in Bassoon-deficient mice. Elife 9 (2020).

83. Montenegro-Venegas, C. et al. Bassoon inhibits proteasome activity via interaction with PSMB4. Cell Mol Life Sci 78, 1545–1563 (2021).

84. Annamneedi, A. et al. Ablation of the presynaptic organizer Bassoon in excitatory neurons retards dentate gyrus maturation and enhances learning performance. Brain structure & function 223, 3423–3445 (2018).

85. Schattling, B. et al. Bassoon proteinopathy drives neurodegeneration in multiple sclerosis. Nat Neurosci 22, 887–896 (2019).

86. tom Dieck, S., et al. Direct visualization of newly synthesized target proteins in situ. Nat Methods 12, 411–414 (2015).

87. Cho, I. & Chang, J.B. Simultaneous expansion microscopy imaging of proteins and mRNAs via dual-ExM. Sci Rep 12, 3360 (2022).

88. Konagaya, Y. et al. A Highly Sensitive FRET Biosensor for AMPK Exhibits Heterogeneous AMPK Responses among Cells and Organs. Cell Rep 21, 2628–2638 (2017).

89. Schindelin, J., et al. Fiji: an open-source platform for biological-image analysis. Nat Methods 9, 676–682 (2012).

90. Burrone, J., Li, Z. & Murthy, V.N. Studying vesicle cycling in presynaptic terminals using the genetically encoded probe synaptopHluorin. Nat Protoc 1, 2970–2978 (2006).

